# Symmetry in levels of axon-axon homophilic adhesion establishes topography in the corpus callosum and development of connectivity between brain hemispheres

**DOI:** 10.1101/2024.03.28.587108

**Authors:** Alexandros Poulopoulos, Patrick Davis, Cheryl Brandenburg, Yasuhiro Itoh, Maria J. Galazo, Luciano C. Greig, Andrea J. Romanowski, Bogdan Budnik, Jeffrey D. Macklis

**Affiliations:** Department of Stem Cell and Regenerative Biology, and Center for Brain Science, Harvard University, Cambridge, MA, USA; University of Maryland School of Medicine, Department of Pharmacology, Baltimore, MD, USA; Harvard Center for Mass Spectrometry, Harvard University, Cambridge, MA, USA

## Abstract

Specific and highly diverse connectivity between functionally specialized regions of the nervous system is controlled at multiple scales, from anatomically organized connectivity following macroscopic axon tracts to individual axon target-finding and synapse formation. Identifying mechanisms that enable entire subpopulations of related neurons to project their axons with regional specificity within stereotyped tracts to form appropriate long-range connectivity is key to understanding brain development, organization, and function. Here, we investigate how axons of the cerebral cortex form precise connections between the two cortical hemispheres via the corpus callosum. We identify topographic principles of the developing trans-hemispheric callosal tract that emerge through intrinsic guidance executed by growing axons in the corpus callosum within the first postnatal week in mice. Using micro-transplantation of regionally distinct neurons, subtype-specific growth cone purification, subcellular proteomics, and in utero gene manipulation, we investigate guidance mechanisms of transhemispheric axons. We find that adhesion molecule levels instruct tract topography and target field guidance. We propose a model in which transcallosal axons in the developing brain perform a “handshake” that is guided through co-fasciculation with symmetric contralateral axons, resulting in the stereotyped homotopic connectivity between the brain’s hemispheres.

## INTRODUCTION

The mammalian cerebral cortex is “wired” by axon contacts that produce connectivity for sensory integration, high-level dexterous motor control, associative behavior, and higher level cognition. Connectivity in cortex combines recurrent local connections with structured and precise long-range projections for integrative-associative function.^1^ Much of cortical functional specialization stems from topography of long-range connections, established in large part by axon growth cones forming projections from an immense variety of distinct projection neuron subtypes.^2^ The dramatic evolutionary expansion of axonal “white matter” (heavily myelinated) tracts, and the long-range axon projections they contain, highlight the critical roles of long-range projection circuitry in brain function.^3,4^

The largest axonal white matter tract in mammals is the corpus callosum, evolutionarily new in placental mammals, and containing hundreds of millions of axons precisely and bi-directionally connecting homotopic areas of the cerebral hemispheres.^5–8^ Agenesis and dysgenesis of the corpus callosum are associated with substantial disruptions of normal social and cognitive function,^9^ while subtle perturbations in callosal structure and activity are associated with a wide variety of circuit pathologies, including schizophrenia and autism.^10,11^

The early organization and developmental principles of axon projection and target guidance,^6,12^ are only sparsely understood.^13^ Beyond initial development, activity-dependent refinement through sprouting, plasticity, and elimination plays critical roles in maturation of functional intracortical circuitry.^14^ A central question is how growing axons of a projection-neuron subtype select topographically distinct targets during axon growth.

Here, we investigated developmental principles of the nascent corpus callosum during the first postnatal week in mice. We focus on cellular mechanisms guiding callosal axons along their projection course to their correct contralateral target fields. We identify intrinsic axon properties that, through self-organized interactions with peer axons, result in emergence of initial topography of developing interhemispheric callosal projection neuron circuitry.

## RESULTS

### Developing cortex forms trans-hemispheric projections with an early, spatially continuous corticotopic map

We first investigated trans-hemispheric projection patterns of the mouse cerebral cortex during early postnatal development. We modified a lipophilic dye tracing approach to map axon projections relative to their point-of-origin on the surface of the cortex. We implanted three nitrocellulose filter squares (approx. 1-2 mm^2^), each pre-loaded with either red, green, or blue (RGB) lipophilic dye, placed along the surface of the cortex in early postnatal (P1-2) mouse pups in either medial-lateral (ML) or anterior-posterior (AP) orientation. Brains were sectioned and imaged at P4, allowing sufficient time for fluorescent labeling of actively growing axon projections. This diffusion-based approach resulted in remarkably homogenous intensity of labeling of cortical projections, including callosal axons. These mesoscale RGB topographic projection maps reveal the developmental projection patterns along the two neuroanatomical axes of placement (ML, AP; Figures 1 and S1).

**Figure 1.**
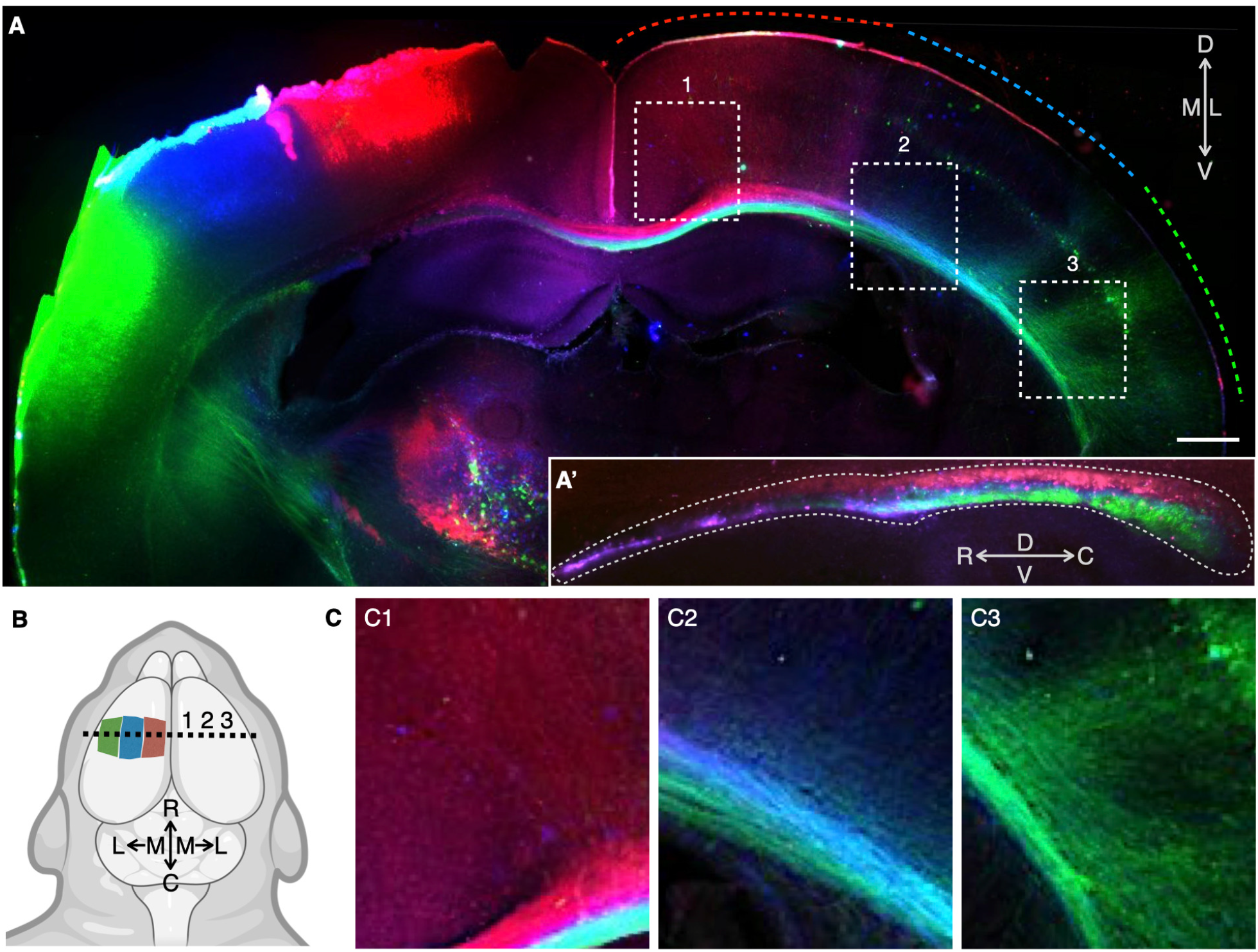
Topographic mapping of cortical projections reveals continuous corticotopic organization of the developing corpus callosum. (A) Coronal section of a P4 brain is shown with continuous mediolateral-axis labeling using neuroanatomical tracers in three colors placed via small, saturated filter paper squares on the cortical surface along topographic axes in infant mice to reveal the internal topography along white matter tracts. Dashed color lines at cortical surface on the right (contralateral) side indicate the approximate coverage of callosal axon termination fields in the contralateral cortex. Areas in numbered dashed boxes are magnified in (C). Inset (A’) shows a sagittal section of the corpus callosum (outlined with dashed perimeter) at the level of the midline. Callosal axon projections (projecting orthogonal to the page) display corticotopic topography throughout the callosal white matter tract such that the mediolateral axis in the cortex is represented by the dorsoventral axis in the callosum. D, dorsal; V, ventral; M, medial; L, lateral; R, rostral; C, caudal. Scale bar corresponds to 500 μm in the main panel, 260 μm in the inset (A’), and 180 μm in panels (C). (B) Schema of mediolateral-axis labeling used in (A). Anatomical axes labeled in (A) are schematized. Dashed like shows the coronal sectioning-plane of the main panel in (A). Numbers show relative positions of the numbered, dashed boxes in (A), magnified in (C). Antero-posterior axis labeling is shown in Figure S1. (C) Labeled transhemispheric axons in the corpus callosum and grey matter of the contralateral hemisphere numbered corresponding to the numbered dashed boxes in (A).

These first-level investigations reinforce that, as previously described,^6,8,15^ the dominant topographic feature of callosal projections is their “homotopic” map. Most trans-hemispheric axons form a symmetric projection path by targeting the mirror-image positions of their parent cell bodies in the contralateral cortex. This homotopic map is evident in both the ML (Figure 1) and AP axes (Figure S1), indicating it manifests as a spatially continuous corticotopic map along the entire mapped extent of the cortex as early as P4.

We assessed RGB labeling beyond the first postnatal week, revealing that mature callosal projections maintain their homotopic mapping throughout cortex. However, the spatial continuity of the map is interrupted, with distinct columns of dense homotopic innervation interspersed with areas largely devoid of callosal axons.^16^ This is consistent with activity-dependent pruning beginning in the second postnatal week, resulting in the mature discontinuous homotopic map characterized by callosal and acallosal cortical areas.^17–19^ Together, these results indicate that callosal axons develop a spatially continuous, symmetric topographic map of the cortex, then prune into mature patches of area-specific homotopic connections, as is observed in the human brain.^15^

The data reveal strikingly topographic patterning, evident throughout the internal organization of the callosal tract. This is present well before axons approach their target fields (Figure 1, inset). Axons emanating from neighboring neuron cell bodies within the same color patch of the RGB tracing system remain neighbors throughout their growth across the callosal tract. The relative cell body positions of cortical grey matter are represented in the relative axon positions in the developing callosal white matter; the spatially continuous ML axis in grey matter corresponds to the dorsoventral (DV) axis in callosal white matter (Figure 1), while the AP axis is maintained as spatially continuous in both (Figure S1). This simple principle comprises the dominant features of the early corpus callosum: homotopic innervation of the target field and corticotopic organization of the tract.

Interestingly, though homotopic projections are by far the dominant feature of the callosum, our mesoscale data reveal evidence of small subsets of axons that do not project to homotopic target fields (e.g. callosal axons targeting the claustrum in Figure 1A), consistent with previous reports.^12^ Our data also reveal a small subset of axons that stratify non-topographically within the callosal tract. This minority subset of axons originate from a mix of areas in the cortex, represented by labeling in all three RGB channels, and appear to segregate from corticotopic axons in the white matter to form a thin layer of non-corticotopic, mixed-color axons at the rostral and ventral fringe of the midline callosum (Figure 1D).

Together, these data identify that the vast majority of interhemispheric axons maintain spatially continuous, “neighborly” relations with other axons throughout their projection trajectories. These relations are based on the corticotopic position of their cell bodies along the ML and AP axes of cortical grey matter. This is reflected both in the resulting homotopic target fields of axons in contralateral cortex, as well as in the internal corticotopic organization of axons in the callosal white matter. These data suggest that intracortical axons execute a perinatal growth and guidance program that determines their internal position in the white matter tract, and their eventual exit position in their contralateral target fields to form function-specific circuitry.

We next investigated whether developing callosal axons maintain this striking corticotopic organization as they grow toward their target areas, similar to pre-target sorting reported for sensory axon projections.^20–22^ We modified an *in utero* electroporation approach^23^ to label superficial-layer callosal projection neurons from distinct cortical areas. By separately injecting GFP and RFP plasmids into the two lateral ventricles, and by carefully adjusting the orientation of the electroporation electric field via positive and negative paddle positions, we effected dual, bilateral labeling with distinct colors, differentiating medial versus lateral callosal projection neurons in the same brain (Figure S2).

This labeling approach confirmed the internal corticotopic organization of the corpus callosum, revealing a conversion of ML position in grey matter to DV topography within the callosal tract itself (Figure S2). This labeling further revealed that, as early as P2, medial and lateral axons with growth cones still in the callosal white matter are already sorted, well before entering their grey matter target fields, thus confirming pre-target sorting in the callosal tract (Figure S2A). Together, these data indicate that developing callosal tract axons grow in close proximity to axons of neighboring callosal projection neurons during the first postnatal week, forming a relatively continuous topographic map throughout the developing callosal tract, and connecting to homotopic target fields on the contralateral cortex.

### Callosal axons intrinsically project to homotopic target fields

At least two broad mechanistic models might explain the development of the corticotopic patterns revealed by mesoscale projection mapping. In an “extrinsic” model, the substratum of the nascent white matter tract has predefined topographic features that growing callosal axons passively follow with simple forward elongation. In this case, the substratum would sort and guide growing axons toward their contralateral homotopic target fields based on the ipsilateral position of entry into the callosum. According to this model, callosal growth cones would provide only the directional growth toward the midline and beyond to the contralateral cortex, while “agnostic” to the specific position of their target fields. These axons would follow the extrinsic substrate in a sense as “train tracks” guiding growth cones along prepatterned homotopic trajectories. This mechanism has been proposed as the primary determinant of homotopic projection in the corpus callosum.^12^

In an alternative “intrinsic” model, callosal axons from distinct areas of the cortex are inherently distinct (via molecules and/or activity), such that they execute intrinsic guidance toward their destination target areas, collectively producing the homotopic topography that emerges during early development of the callosal tract.

We devised an experiment that would distinguish between these “extrinsic” and “intrinsic” models of callosal development (Figure 2A). Callosal projection neurons from lateral cortex could be extracted and ectopically transplanted to a position in medial cortex. The ectopic neurons would be allowed to grow their axons alongside native callosal axons from the host medial cortex.

**Figure 2.**
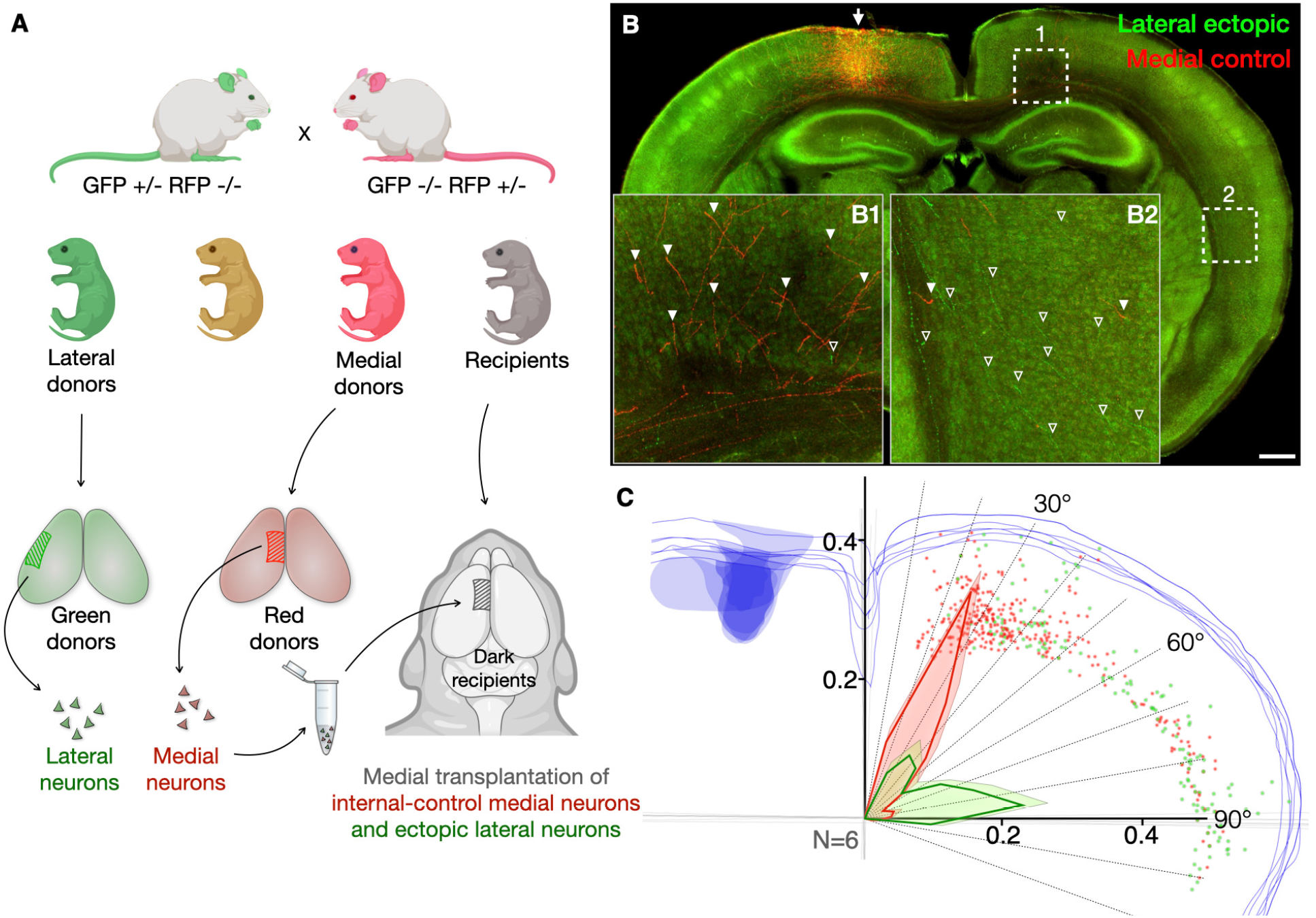
Heterotopic micro-transplantation of lateral and medial cortical neurons reveals intrinsic callosal axon targeting. (A) Schematic of the heterotopic and simultaneous control syntopic micro-transplantation experimental design. GFP +/- mice were bred with RFP +/- mice. The two loci are independent, resulting in Mendelian 25% frequencies of red, green, yellow, or dark F1 offspring. Neural micro-transplantations were performed within the first 6 h after birth. Cells from lateral cortex of green littermate donors were mixed with cells from medial cortex from red littermate donors, and micro-injected into medial cortex of dark littermate recipients. Recipient brains were imaged for GFP and RFP at P7. (B) Recipient dark brain at P7 (non-specific green glow is autofluorescence that helps to contour neuroanatomical features) hosting both ectopic transplanted cells from lateral cortex (green) and internal-control transplanted cells from medial cortex (red) together micro-injected into medial cortex of dark recipients. Arrow points to micro-injection site. Insets B1 and B2 show enlarged fields from dashed boxes from (1) medial and (2) lateral contralateral cortex. Axons from transplanted neurons. Many red axons are project medially (arrowheads in B1), very few laterally (arrowheads in B2). In contrast, many bright green axons are visible above the autofluorescence projecting laterally (open arrowheads in B2), with only a few medially (open arrowheads in B1). Scale bar corresponds to 500 μm in the main panel and 100 μm in insets (B1 and B2). (C) Quantification of medial-derived red control and lateral-derived green axon positions in contralateral grey matter. Traced schematic of brain sections from one transplant-recipient mouse brain. Blue outlines coronal sections and transplantation site. Red and green dots indicate corresponding axon termination positions. Grey lines indicate midline and meridian axes used to quantify axon position as degrees from midline at intersection. See Figure S3 for equivalent tracing overlaying sections from all 6 transplant-recipient brains showing transplantation site and targeting consistency. Angle histogram with 10º bins (dashed angled lines) summarizes measurements from 6 transplant-recipient mice. Heavy green and red trend lines show the proportion of corresponding axons in each bin, while the thin enveloped lines show mean + standard error in each bin. The major fraction of green axons (0.59) from lateral-derived neurons intrinsically target lateral cortex (60º-100º binned range), where only a minor fraction (0.12) of control red axons from medial-derived neurons target. The major fraction of red axons (0.79) target medial cortex (10º-50º binned range), where only a minor fraction (0.29) of green axons target.

The “extrinsic” model (“train tracks”) predicts that axons from lateral-derived neurons would enter the callosal tract in the medial cortex and grow following the trajectory of a prepatterned substratum toward the medial homotopic area on the contralateral cortex. In this model, callosal axons across the cortex are all equivalent and “agnostic” of their target fields. As such, the ectopic lateral callosal axons would behave like natively medial callosal axons and follow the medial-to-medial “train tracks” to target positions in the contralateral homotopic medial cortex.

By contrast, the “intrinsic” model predicts that the ectopic neurons would intrinsically display a tendency to project their callosal axons to lateral contralateral cortex, even though they now project from cell bodies in medial cortex. Such medial-to-lateral (heterotopic) projection, contrary to native medial-to-medial (homotopic) projections from the medial cortex, would indicate that developing axons project with native molecular tropism toward their originally appropriate target fields, and that axons themselves possess intrinsic topographic information for positioning in the nascent white matter tract.

We tested these two alternative models using a transplantation approach in neonatal mouse brain. We micro-transplanted genetically green fluorescent neurons from lateral cortex into medial cortex along with medial neurons of a genetically red mouse, and measured the target field positions of their axons in contralateral cortex. This internal control of co-transplaned neurons from medial cortex both controls for the procedure itself and enables quantitative assessment of targeting. Specifically, ratiometric measurement of projection topography quantitatively assesses the contribution of the intrinsic and extrinsic models in callosal development.

We isolated cortical neurons at P0 from lateral cortex of green (GFP-expressing) transgenic mice, and cortical neurons from medial cortex of red (RFP-expressing) transgenic mice. We combined equal numbers of green/lateral and red/medial neurons in a single cell suspension, and injected into the medial cortex of non-fluorescent mice. For this micro-transplantation experiment, the GFP and RFP donor mice, as well as the non-fluorescent recipient mice, were all newborn littermates produced by heterozygous GFP+/- x RFP+/- breeding to produce litters that include red, green, and non-fluoresent (“dark”) pups (Figure 2A). In this way, micro-transplantation is isogenic and isochronic between test, control, and recipient cells, to minimize potential confounding effects of genetic background and/ or developmental timing on experimental outcomes. Following the micro-transplantation procedure, medial cortex of dark pups hosts green neurons from lateral littermate cortex (ectopic test neurons), and of control red neurons from medial littermate cortex (syntopic control neurons). Transplanted recipient mice were perfused at P7, and brains were prepared and imaged to quantify the target positions of transcallosal green test and red control axons of transplanted neurons.

We aimed to quantify the degree to which projection to specific target fields is intrinsic to growing axons, versus the degree to which axons passively grow on a substrate that extrinsically determines projection target depending on the area from which axons project. These two “limit models” represent the boundaries of a spectrum between cell-autonomous and non-cell-autonomous targeting mechanisms. Red internal control axons serve as a reference for potential effects of transplantation itself. Thus, deviations of targeting by green axons from that of red axons are due to intrinsic properties of axons projected by neurons transplanted from lateral cortex.

These experiments reveal that neurons derived from lateral donor cortex (green) and transplanted into medial host cortex project overwhelmingly to lateral cortex, reflecting intrinsic site-of-origin targeting rather than targeting that is native to their new host area. In striking contrast, control neurons derived from medial cortex (red) and transplanted into medial host cortex project predominantly to medial cortex, reproducing the endogenous homotopic map (Figure 2). These quantitative data indicate largely cell-autonomous mechanisms, with axons executing their intrinsic projection path, which is already determined by P0.

### Electrical Activity does not instruct early targeting of callosal projections

These results indicate that callosal projection neurons are already committed at P0 to project to appropriate target locations, and that they possess information that directs axons to their target fields during the first postnatal week. We next investigated whether this intrinsic information is related to electrical activity patterns known to emerge in upper layer cortical neurons during early postnatal development.^24,25^ We employed a similar experimental approach to address this question with similar ratiometric quantification and internal controls. We employed a new genetic mosaic plasmid system we recently developed^26^, termed BEAM (Binary Expression Aleatory Mosaic), which enables by a Cre-dependent Cre amplification and delay feature quantitatively controllable genetic manipulation of a subtype-specific neuron population, with two distinct genetic populations spatially interdigitated in the same developing brain. We applied BEAM via *in utero* electroporation. We modified half of the superficial layer callosal neurons to become red wild-type control neurons, and the other half to become green test neurons . We achieved roughly 50:50 ratios by titration of initial applied Cre such that half of the electroporated neurons amplify Cre and resultingly express Cre-on GFP together with the test payload, while the other half do not receive Cre, and thus express Cre-off RFP. These RFP-labeled neurons serve as internal controls in the same region and subtype (Figure 3A).

**Figure 3.**
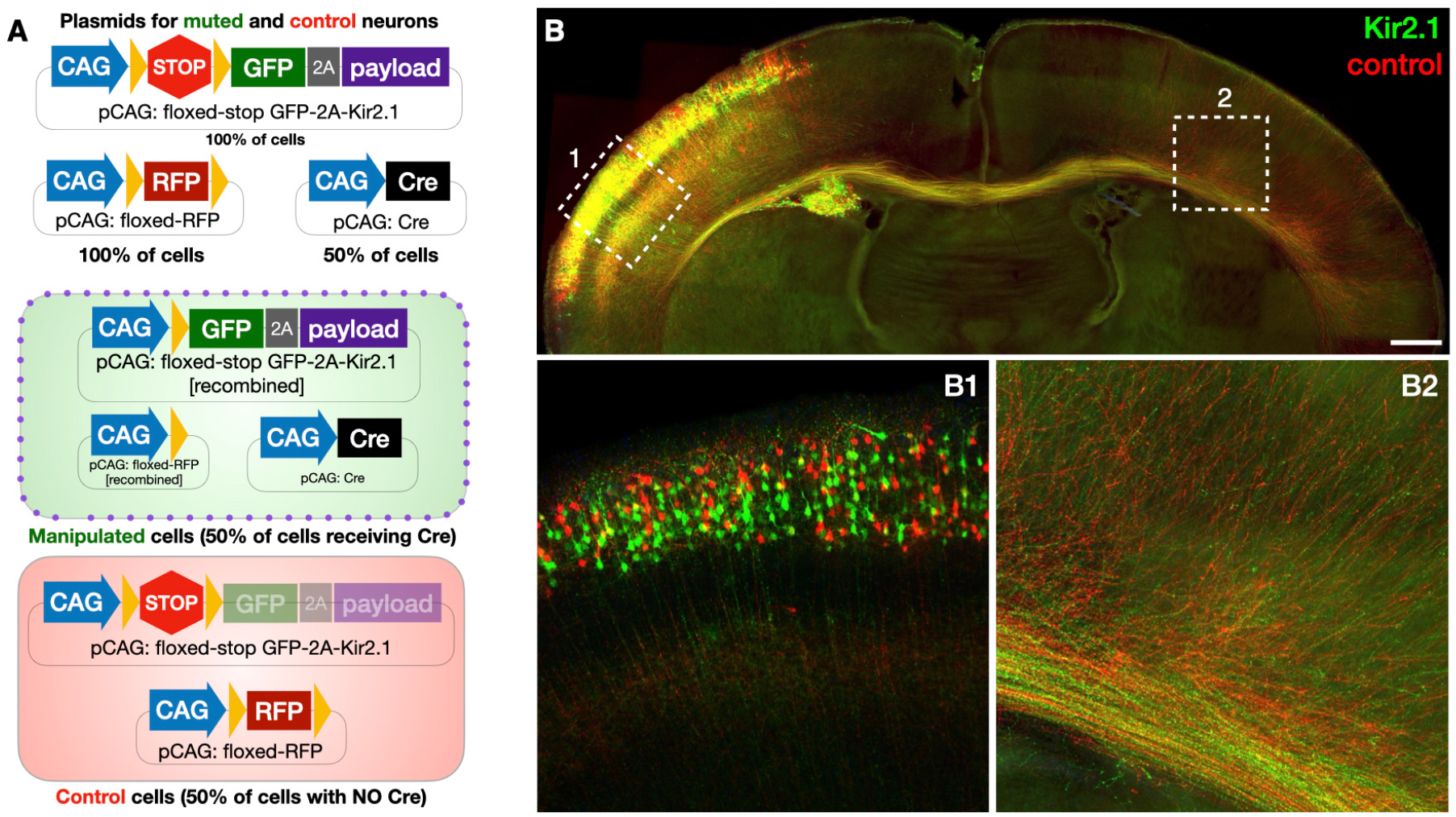
*In utero* genetic mosaic manipulation with spatially co-localized, interspersed internal control neurons (via BEAM^26^) reveals that activity is dispensable for early callosal projection topography. (A) Schematics of the multi-construct, genetic mosaic plasmid system (binary expression aleatory mosaic; BEAM^26^) and the resulting binary expression outcomes in electroporated neurons. The three plasmids are titrated so that only roughly half of all electroporated cells receive a Cre expression plasmid, while all electroporated cells receive a Cre-on manipulation plasmid conditionally expressing GFP and the payload gene (Kir2.1, in this experiment) and a control plasmid expressing RFP only in the absence of Cre. Neurons receiving all 3 plasmids become green manipulated neurons, while neurons receiving only 2 plasmids are red control neurons. (B) Coronal brain section at P7 with mosaic *in utero* electroporation of superficial layer callosal projection neurons stochastically expressing either green or red plasmids (inset B1), interspersed within the same electroporation field (∼2-5% are yellow dual-expressing neurons^26^). Green “experimental” neurons express GFP together with the inward rectifying K^+^ channel Kir2.1 to mute electrical activity. Red “control” neurons express RFP alone, and serve as region- and subtype-specific, spatially interspersed internal controls exhibiting normal activity. Magnified insets of dashed boxed areas of (1) binary experimental-green and control-red neuronal somata in the electroporation field and (2) similarly binary axons at the white matter exit field in contralateral cortex. Scale bar corresponds to 500 μm in the main panel and 100 μm in the insets.

We used BEAM to express the inward rectifying potassium channel (Kir2.1) in GFP+ neurons at E15. Expression of Kir2.1 induces hyperpolarization and disruption of native activity in cortical neurons.^17,19^ These experiments revealed that Kir2.1 expression had no effect on the targeting of GFP+ callosal projections compared to red control axons at P7 (Figure 3B), indicating that electrical activity does not instruct the selection of target field during the first postnatal week, when callosal projections form. Confirming previous reports, we observed disruption of callosal projection maturation after the second postnatal week, consistent with the onset of activity-dependent pruning and refinement of the initial callosal projections.^17^ These data indicate that intrinsic molecular constituents in axons, and not electrical activity, determine projection target fields during early postnatal development of the cerebral cortex.

### Canonical axon guidance molecules do not instruct topographic projection of callosal axons

We next hypothesized that projection target information might be encoded through differential expression of known axon guidance molecules^27,28^ expressed by callosal projection neurons and present in their axons.^12,29^ We tested this hypothesis with three known axon guidance molecules that we previously determined are highly expressed by perinatal superficial layer callosal projection neurons.^30^ We used BEAM at E15 to deliver Cre with Cre-ON GFP and Cre-OFF control RFP into superficial layer cortical projection neurons in three conditional mouse-lines with homozygous floxed alleles for Ephrin-A4 (*Efna4*^flox/flox^), Plexin-D1 (*Plxnd1*^flox/flox^), and Neuropilin-1 (*Nrp1*^flox/flox^). In these mice, green neurons have the floxed gene deleted, and red neurons are internal controls. When imaged at P7, gene deletion of *Plxnd1, Efna4*, or *Nrp1* had no effect compared to internal controls on the tract topography or the target field of the nascent callosal projection (Figure S4). One candidate, the surface semaphorin receptor Neuropilin-1, warrants further comment. A previous study reported that *Nrp1* deletion via electroporation perturbed intra-callosal topography.^12^ Using BEAM, we confirm that, while intra-callosal topography is affected by *Nrp1* KO ipsilaterally, final homotopic projection mapping is unperturbed, indicating a rectifying mechanism after midline-crossing. Our later experiments are informed by this intriguing result.

### Two-color growth cone proteomics identifies topography-associated differences in levels of adhesion molecule Ncam1

We next investigated potential differences in the local proteomes of callosal axon growth cones that derive from topographically distinct areas of cortex. We postulated that growth cones of medial and lateral callosal axons might have quantitative differences in their local proteomes that enable them to execute projection to their correct, respective homotopic target fields. Such subcellular proteome differences might reflect intrinsic factors that enable callosal axons to execute correct targeting cell autonomously, as revealed in the micro-transplantation experiments (Figure 2).

We previously developed “fluorescent small particle sorting” (FSPS) as an approach to obtain projection-specific subcellular transcriptome and proteome data from growth cones of the developing mouse brain.^23^ Here, we applied this approach with asymmetric, bilateral electroporations (Figure S2) to perform two-color differential FSPS of growth cones of medial versus lateral upper-layer callosal projection neurons differentially labeled with distinct fluorescent proteins (Figure 4A). At P2, we dissected cortices, isolated growth cones bilaterally, and performed FSPS to separate and purify medial (red) and lateral (green) callosal growth cones (Figure 4B). We isolated protein and analyzed the two callosal growth cone samples with label-free quantitative mass-spectrometry (LFQ-MS) to compare medial vs. lateral callosal growth cone proteomes.

**Figure 4.**
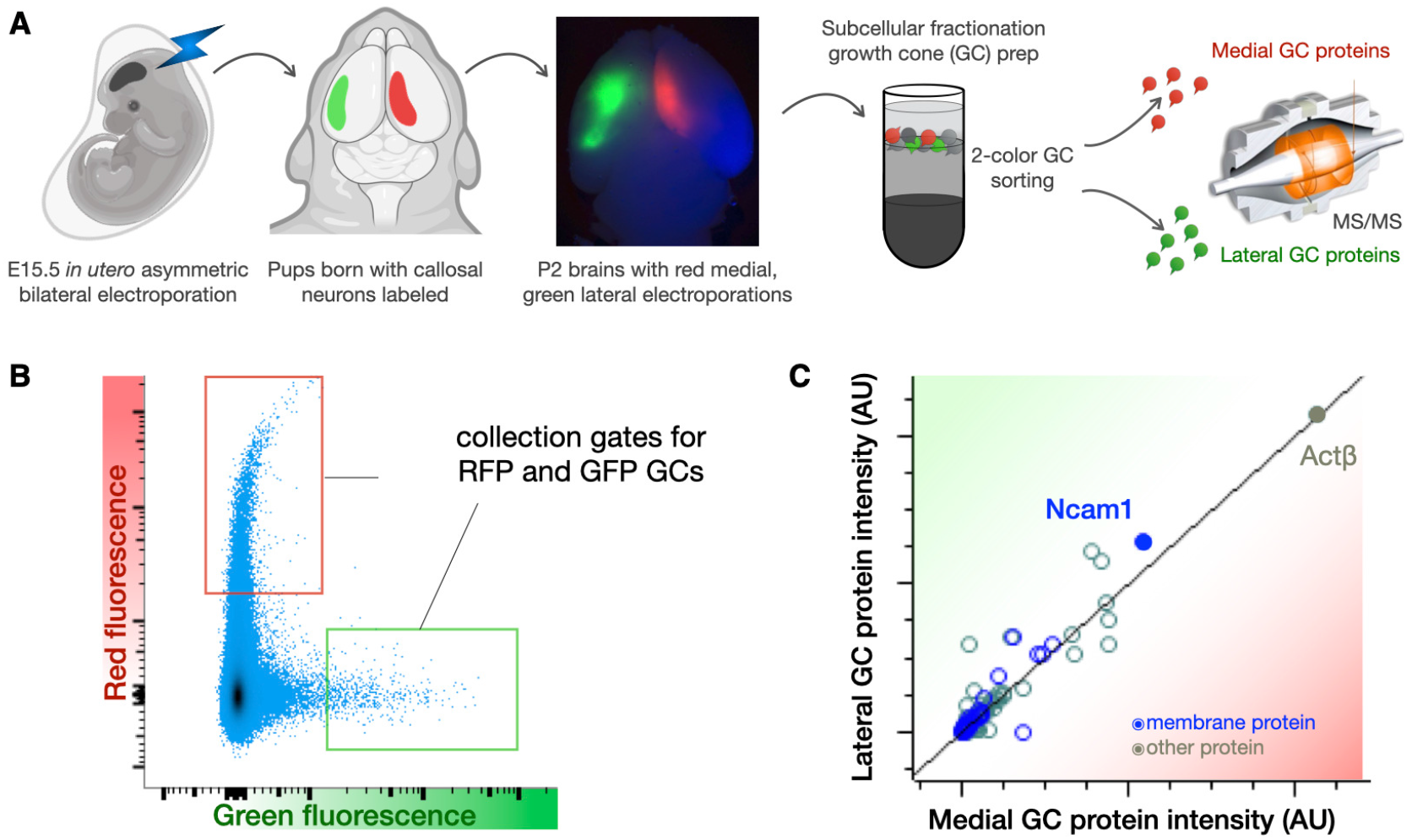
Two-color growth cone fluorescence sorting from medial vs. lateral callosal axons. (A) Schematic of dual asymmetric electroporation to label medial (red) vs. lateral (green) superficial layer callosal projection neurons, subcellular fractionation to isolate growth cones, two-color growth cone purification by fluorescence small particle sorting (FSPS), and label-free proteomics to identify differential protein abundances in medial vs. lateral callosal growth cones. (B) Fluorescent small particle sorting (FSPS) plot of two-color growth cone (GC) sorting. Collection gates for red and green growth cones are indicated. (C) Scatter plot of protein amounts in distinct medial vs. lateral growth cone proteomes (for uncropped axis range, see Figure S5). Each circle represents the average protein intensity measurements from medial (x-axis) and lateral (y-axis) growth cones from two biological replicates, each from 3-5 pooled dual-electroporated brains. Membrane proteins are indicated in blue. Neural cell adhesion molecule 1 (Ncam1) and β-actin (Actβ) are indicated with solid circles. Ncam1 is the highest intensity membrane protein in GCs, and is present at higher levels in lateral GCs, while β-Actin is present at equal levels in both medial and lateral GCs, serving as a quantitative control.

Medial and lateral callosal growth cones contained the same proteins overall, as would be expected from axons of the same overall subtype with shared elements of initial midline directionality, midline crossing, and cortical innervation. We next compared the relative amounts of each protein in the comparative growth cone samples (medial, lateral) by plotting the average LFQ-MS values (Figures 4C). Those high-abundance proteins ubiquitous to growth cone machinery, such as β-Actin, displayed equal intensities between in the two populations. Low abundance proteins displayed higher variation between the two samples, as expected from low signal-to-noise ratios in these ranges (Figure 4C; represented in the spread at the base of the scatter plot).

We hypothesized that differences in levels of specific proteins might differentially instruct callosal axon topography. We reasoned that cell surface proteins are biologically likely candidates for such a role, and we considered differences appearing in proteins of higher abundance as quantitatively more reliable. With these considerations in mind, we filtered the subcellular proteome data for cell surface proteins that deviate from the equimolar diagonal to select candidate proteins putatively present at different levels in medial vs. lateral callosal growth cones. These candidates were functionally investigated individually using BEAM for their influence over targeting based on their expression levels.

By far the most abundant cell surface protein displaying deviation in protein level between medial and lateral callosal growth cones is the homophilic adhesion molecule Ncam1. It has approximately 20% higher levels in lateral vs. medial growth cones. We thus investigated whether over-expression of Ncam1 would shift transcallosal targeting laterally. We employed BEAM to produce spatially interspersed, genetically mosaic populations of Ncam1 over-expressing neurons and control neurons with unmanipulated Ncam1.

Quite strikingly, Ncam1 overexpressing axons projecting from cell bodies in the electroporation field progressively separated from internal control axons, both within the corpus callosum itself, and once near contralateral target fields. Ncam1 overexpressing axons traversed the callosum ventral to their control peers (Figure 5A and Figure S6), consistent with the normal midline topography of laterally projecting callosal axons (Figure 1). Contralaterally, Ncam1 overexpressing axons preferentially exited the callosum to innervate target fields in the far lateral cortex (Figure 5A3) but not the more medial, homotopic somatosensory area to which control axons project (Figure 5A2).

**Figure 5.**
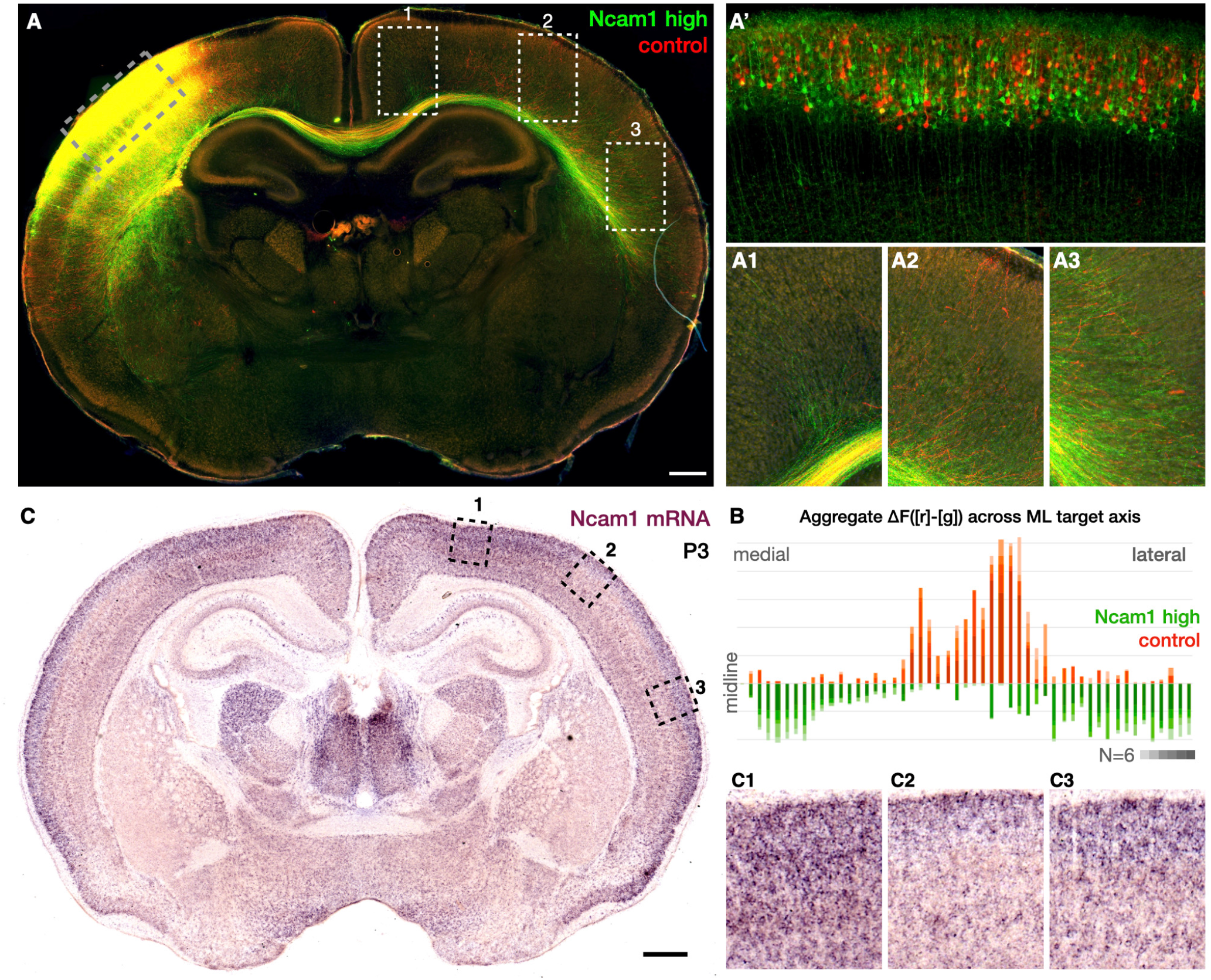
Ncam1 expression levels determine axon targeting of callosal projection axons. (A) Genetic mosaic *in utero* electroporation of superficial layer callosal projection neurons. Green electroporated neurons over-express Ncam1 (“Ncam1 high”) compared to red electroporated wild-type control neurons (“control”). Coronal overview at P7 with magnified insets from dashed boxes: green Ncam1 high and red control electroporated neuron cell bodies reside in somatosensory cortex; (A’) Ncam1 and control neurons interspersed in the same spatial domain; (A”) axons from interspersed Ncam1 high (green) and control (red) cell bodies already substantially segregated at the callosal midline; (A1, A3) Ncam1 high (green) callosal axons target non-homotopic areas of contralateral cortex that are medial (A1) and lateral (A3) to the homotopic somatosensory area, which is targeted by control (red) axons (A2). Scale bar corresponds to 500 μm in the main panel, 60 μm in inset (A’), and 180 μm in insets (A1-A3). (B) Quantification of Ncam1 high (green) vs. control (red) axon targeting in contralateral cortex. Plot shows 50 bins dividing the mediolateral axis of the contralateral hemisphere, beginning at the midline, running along the deepest layer (6B) of cortical grey matter, and ending at the claustrum. Within each sample, green (Ncam1 high) fluorescence intensity is subtracted from red (control) fluorescence intensity. ΔF([r]-[g]) is plotted to display the genotype bias of axon targeting within that bin. Positive values denote red (control) preferred axon targeting, and negative values denote green (Ncam1 high) preferred axon targeting. Values from N=6 electroporated mice are aggregated in each bin, separated by color opacity. Horizontal lines represent increments of 10% normalized intensity difference ΔF. (C) *In situ* hybridization for Ncam1 expression in a coronal section at P3. Magnified insets C1 to C3 of areas from medial to lateral cortex corresponding to axon target fields in insets A1 to A3. Green (Ncam1 high) axons target cortical areas A1 and A3 that have endogenously high Ncam1 expression (C1 & C3), while avoiding the homotopic area of moderate Ncam1 expression (C2) endogenously targeted by red (control) axons (A2). Scale bar corresponds to 500 μm in the main panel and 120 μm in the insets (C1-C3).

Unexpectedly, a subset of Ncam1 overexpressing axons project to the far medial cingulate area, another area to which control axons do not project. We thus examined the endogenous expression levels of Ncam1 along the mediolateral topographic axis of P3 brains using mRNA *in situ* hybridization. We identify graded expression of Ncam1 in superficial-layer cortex characterized by high levels in far lateral and far medial areas, and lower levels in the intermediate area (Figure 5 B1-3).

This graded pattern of Ncam1 expression mirrors the projection pattern revealed by the BEAM experiments. Axons overexpressing Ncam1 preferentially target non-homotopic areas that endogenously express high levels of Ncam1. Together, these experiments reveal both that Ncam1 expression differs across cortical areas, and that Ncam1 protein levels similarly differ in growth cones of axons across the same cortical areas. Further, manipulation of Ncam1 levels modifies axon targeting toward alignment of Ncam1 levels with the endogenous levels of the target area.

### Model of axon projection by matching of molecular adhesion levels

Taken together, these experiments and results propose a mechanistic model of transhemispheric axon development. Though we will focus here on a single identified molecular component of this model– graded levels of Ncam1– the model is generalized regarding a potentially broad range of axonal homophilic cell adhesion molecules (hCAMs). This model postulates that hCAMs mediate interactions^31^ through which axons recognize and optimally adhere to other axons emanating from their nearest ipsilateral neighboring neurons (*ciscallosal* axon-axon interactions).

This process enables callosal axons to grow toward the midline while maintaining cortical topography (Figure 6A). Upon reaching the midline, callosal axons encounter their time-matched and area-matched counterparts from the contralateral hemisphere. We propose that, from the midline onward, callosal axons additionally engage in hCAM interactions with contralateral callosal axons (*transcallosal* axon-axon interactions). Through recognition of and co-fasciculation with contralateral axons originating from the homotopic contralateral area, transcallosal axons continue growth, achieving both pre-target sorting and final targeting to the homotopic area, from which peer contralateral axons emanate (Figure 6B).

**Figure 6.**
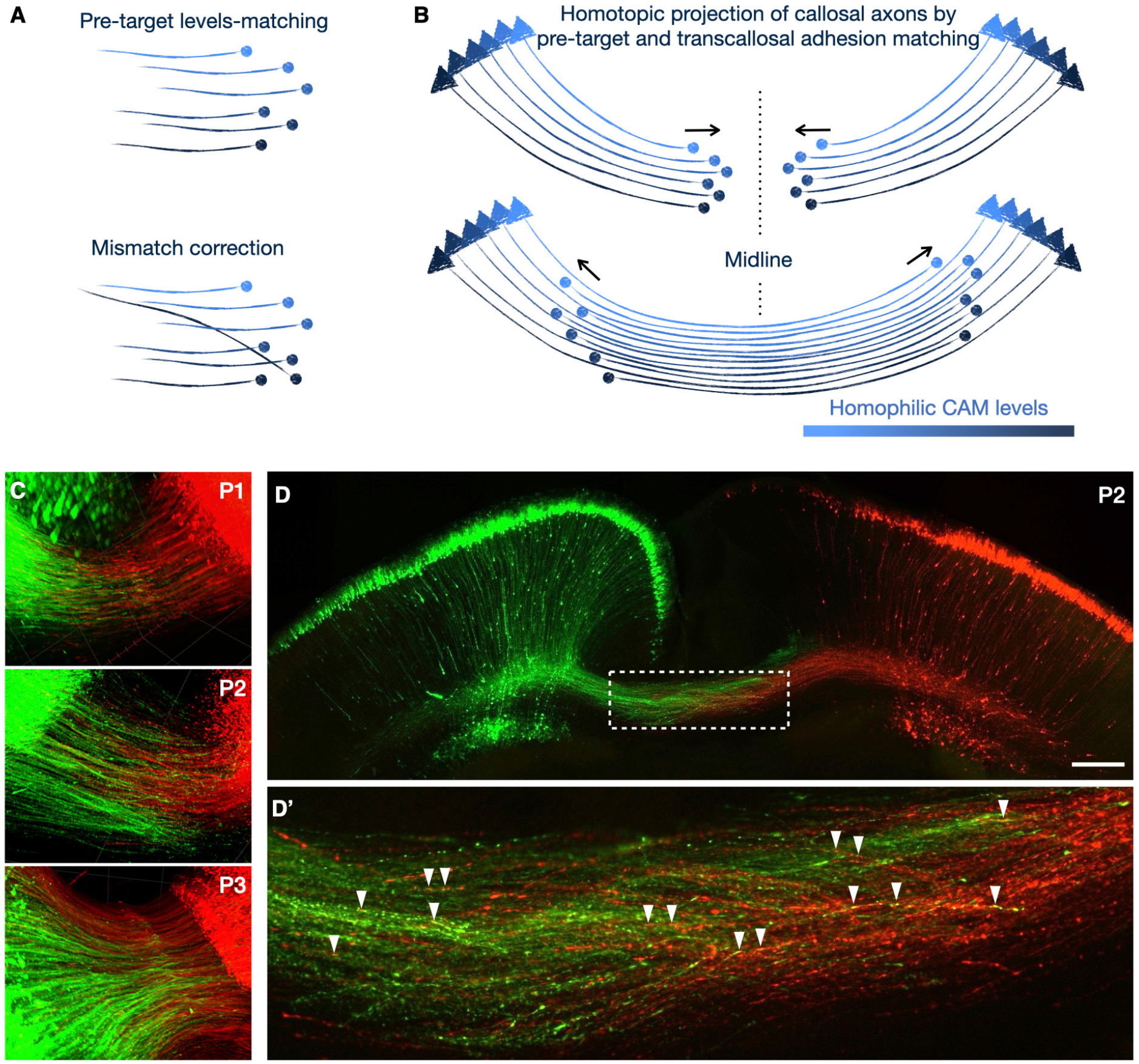
Model of corticotopic targeting by matching levels of homophilic adhesion in callosal axon projections. (A) Schema showing tips of axons with growth cones (round ends) and graded levels of adhesion molecule (indicated by darkness range) maintaining topography as they form a projection. Growing axon tracts display population behavior maximizing the total number of adhesion complexes by axons with similar levels positioning themselves next to each other. Under conditions of topographic expression gradients, axons of neighboring neuron cell bodies have similar expression levels, and thus grow together throughout the tract. This is schematized as pre-target topographic sorting by matching of adhesion levels (left panel). Under conditions of complex discontinuous expression patterns, or in the case of ectopic micro-transplantation (as in experiment of Figure 2), axons with distinct adhesion molecule levels participate in iterative axon-axon interactions and progressively process toward positions that most closely match their own adhesion levels. This is schematized as mismatch correction (right panel). This generalizable model can potentially incorporate multiple or many homophilic adhesion molecule systems, resulting in integrated population behavior to maximize overall adhesion complex occupancy by optimally matching adhesion levels of each component system. (B) Schematic of the model of matching of adhesion levels applied to the bilaterally symmetric development of the corpus callosum. Callosal projection neurons express homophilic adhesion molecules (e.g. Ncam1) in topographic gradients of their cell body positions along the cortex, such that the level on a given axon depends on the position of its cell body (indicated by triangles) along the cortical plane. Callosal axons from each hemisphere grow synchronously toward the midline, maintaining topographic organization via pre-target sorting via matching of axon-axon molecular interaction levels (top panel). Once callosal axons cross the midline, they continue to match adhesion molecule levels via axon-axon interactions. Due to bilateral symmetry of expression gradients, the closest matching levels are present on axons from the mirror image location in the contralateral cortex, resulting in callosal axons targeting the symmetric position in the contralateral hemisphere. (C) Time course of transcallosal axon development from P1 to P3, revealed with light sheet microscopy on cleared brains of symmetric dual-hemisphere electroporation. GFP and RFP label the same callosal neuron populations on separate cortical hemispheres. Their growing axons meet and co-fasciculate as they cross the midline. See Movie S1 for full P2 brain. (D) Coronal section of P2 brain showing callosal axons’ homotopic “handshake” after symmetric dual-hemisphere electroporation. (D’) Inset of dashed box in D shows red and green axons from contralateral hemispheres co-fasciculating after crossing the midline (arrowheads indicate representative positions of close apposition of red and green axons appearing yellow). Individual red and green axons can be seen juxtaposed in close contact over long distances with opposite trajectories within the axon tract, consistent with transcallosal fasciculation. Scale bar corresponds to 250 μm in the main panel and 50 μm in the inset.

This proposed mechanism postulates recognition specificity of *ciscallosal* nearest-neighbors as well as *transcallosal* homotopic contralateral axon-axon interactions. We posit that, in both cases, specificity emerges from population behavior according to the principle of maximizing hCAM binding within the white matter tract. Growing axons will preferentially co-fasciculate with axons that have similar levels of hCAMs (Figure 6A), other of which likely remain to be identified. While two axons with different hCAM levels will interact, they can only form the level of density of adhesion complexes present in the lower-expressing axon, thus leaving a surplus of unengaged hCAMs on the higher-expressing axon. Since neighboring neurons in cortex tend to have similar expression profiles from developmental gradients,^32^ the maximum density of axon-axon adhesion complexes in the corpus callosum would be achieved when axons are arranged corticotopically throughout the white mater tract. Indeed, this is the organization revealed by mesoscale mapping of the callosal projection (Figures 1, S1, and S2).

Moreover, this principle of self-organization of callosal topography is highly consistent with the results from the ectopic micro-transplantation experiments, in which axons from ectopic neurons express hCAMs at different levels from their interspersed neighbors within the shared implantation site. While these axons initially engage in local non-level-matched axon-axon interactions, they progressively shift interactions to more appropriate ipsilateral and contralateral positions within the dorso-ventral– thus medio-lateral targeting– axis of the callosum. As they grow, this model predicts that they would tend to align their dorso-ventral location and eventual direction of growth with axons more closely matching their own levels, gradually positioning themselves within the tract alongside axons from the area from which ectopic neurons originated (Figure 6A). The model thus predicts that, ultimately, transplanted axons will exit the callosum near the area homotopic to their origin, rather than that of their ectopic site of implantation. The experimental data (Figure 2) are strongly consistent with this prediction and this model.

The mechanism we propose predicts the robustly homotopic nature of the transcallosal map, directed by the bilateral symmetry of gene expression in the cerebral hemicortices. Callosal axons are predicted to encounter their contralateral hCAM-matched axons at the midline, with “matching” including expression of an identical repertoire of hCAMs at the same levels. In the respective contralateral domains of the callosal tract, these axons are predicted to engage in transcallosal axon-axon interactions, following each other all the way to their respective contralateral homotopic areas, from which their counterpart axons originated (Figure 6B).

### Transcallosal fasciculation in developing cortical white matter

We directly investigated whether transcallosal fasciculation occurs when axons grow across the midline, as the model predicts. To do so, we devised a strategy to follow time-matched and area matched callosal axons from both hemispheres as they meet at the midline during early postnatal development. We used wide-field tritrode electroporation to label superficial-layer neurons in each hemisphere concurrently, along with bright red or green fluorescent protein variants unique to each hemisphere. We imaged labeled axons in the intact brain with light sheet microscopy after clearing to produce a neonatal time-course between P1 and P3 (Figure 6C). This imaging reveals that labeled callosal axons meet their contralateral counterparts at the midline at P1, and continue to grow interdigitated with axons from the contralateral hemisphere at P2 and P3. Strikingly, labeled axons from opposite hemispheres are revealed to co-fasciculate, even appearing occasionally to track along one another in opposite directions (Figure 6D and Movie S1). These results directly support the occurrence of transcallosal axon-axon interactions, and suggest a function of these interactions in guiding appropriate, area-specific topographic growth in the developing callosum, and thereby in the cerebral cortex.

## DISCUSSION

The spatial organization of white matter tracts both defines the topographic connectivity of the brain and underlies the functional specificity of long-range central nervous system circuitry. The paths axons follow from their cell bodies to their target areas for innervation enable the remarkable precision of circuit connectivity over distances up to ∼10^3^ cell body diameters during development, and up to ∼10^5^ cell body diameters in adulthood. In addition to their characteristic overall trajectories, many tracts exhibit internal topography that organizes how individual axons course through the tract in relation to one another,^28^ as seen in the olfactory nerve,^20^ optic nerve,^21^ pyramidal/corticospinal tract,^33^ hippocampal projections,^34^ and corpus callosum.^12^

Here, we investigated the development of topography within the corpus callosum, the largest white matter tract of the mammalian nervous system, and the foundation of interhemispheric cortico-cortical associative and integrative circuit connectivity. The corpus callosum’s precision underlies appropriate sensori-motor integration between the hemispheres, social-behavioral function,^3511^ and associative connectivity between asymmetrically specialized areas. Our results reveal three developmental features of the early callosum (a-c below), and three mechanistic elements of callosal axon target selection (i-iii below), which together build the early connectivity between the cortical hemispheres with circuit and functional precision.

Our results reveal that: a) the corpus callosum develops as a continuous corticotopic map (Figures 1, S1, and S2); b) areally distinct callosal growth cones contain the same set of proteins, but differ from each other in the levels of some proteins, including Ncam1 (Figures 4); c) callosal axons co-fasciculate with their homotopic contralateral counterparts after crossing the midline (Figure 6 and Movie S1). Early callosal axon target selection is i) executed intrinsically and determined by P0 (Figure 2); ii) not instructed by electrical activity (Figure 3); iii) predictably altered by manipulating the levels of Ncam1 unilaterally (Figures 5 and S6). These results and mechanistic elements integrate into a model of how the brain’s transhemispheric projections are established.

Foundational to the emergent model is that transhemispheric projections form as a continuous “corticotopic” map in which axons assume positions within the callosum according to the positions within the cortex of their parent cell bodies (Figures 1 and S1). This continuous map of cortical areal position during the first postnatal week starkly contrasts with the highly discontinuous homotopic innervation present in the mature cortex, in which characteristic regions, such as primary sensory (S1) and primary motor cortex, are acallosal, while areas such as the S1/2 boundary maintain rich pillars of callosal innervation.^36^ This discontinuous structure emerges in the 2nd postnatal week in mice, during which endogenous activity patterns, dispensable during the first week of callosal development (Figure 3), guide maturation processes that sculpt the continuous callosal map into the mature discontinuous patterns of transhemispheric connectivity via pruning.^19,37^

Axons assume and maintain topographic positions within the callosum as they grow, and already do so before crossing the midline, maintaining this topographical positioning until reaching their target fields (Figure S2). This feature is reminiscent of “pre-target sorting” in olfactory and visual projections, in which axons grow into their positions within the tract based on the olfactory receptor they express,^20^ or based on the location of their cell bodies in the retina,^21^ respectively. Callosal neurons are committed by P0 to intrinsically execute their transhemispheric trajectory based on the position of their cell body in the cortex.

Our experiments directly tested between two alternatives:1) whether position on the cortex is the direct factor guiding axons by way of extrinsic features, such as substrate patterning; or 2) whether location on the cortex translates into intrinsic molecular features of axons that enable individual axons to navigate relatively autonomously to their appropriate corticotopic target area. These experiments reveal that axons are intrinsically committed by P0– even if their somata are heterotopically positioned via transplantation, thus projecting from an ectopic area, they correctly project to their appropriate target area in the contralateral cortex (Figure 2).

It is interesting to note that, when transplantation was performed more than 6 hours after birth, axon projections no longer maintained their topographic specificity. This was true for axons from both control and ectopic neurons. This result indicates that topographic axon projection has a temporal component necessary for fidelity, and suggests a mechanism requiring coordination with dynamic features with in the callosal environment. Our experiments also reveal that callosal axons’ projection to their proper targets does not rely on native electrical activity (Figure 3B) nor on the canonical callosal guidance molecules Neuropilin 1, Plexin D1, or Ephrin A4 (Figure S4).

Rather, callosal axon targeting is identified to be sensitive to the expression level of the homophilic adhesion molecule Ncam1, likely one of multiple such regulators. Exogenous increase of Ncam1 levels results in axons changing their target field from their appropriate homotopic area to areas that intrinsically match the higher levels of Ncam1 expression (Figures 5 and S6). Given the well-characterized function of Ncam1, these results strongly suggest that adhesive axon-axon interactions and co-fasciculation are critically involved in navigation toward appropriate topographic target fields.

Ncam1 is known to mediate axon co-fasciculation in white matter tracts, indicated by the defasciculation phenotypes of Ncam1 gene deletion mice,^38,39,40^ fully consistent with our data indicating its control over matching axons with graded expression levels Interestingly, however, Ncam1 gene deletion mice do not display overt changes in the projection targets, as our model would predict of bilaterally symmetric genetic perturbations, further indicating the broader involvement of other hCAMs in the levels-matching mechanism of axon targeting. Taken together, both sets of results– from prior gene deletion experiments and from our current experiments– support a unifying mechanistic model of callosal axon targeting.

We propose that callosal axons co-fasciculate with their neighbors through Ncam1 and other homophilic adhesion molecules in a manner that pairs axons that share similar expression levels of these molecules. This results in axons from similar areas of the cortex remaining close to their neighbors as they course through the callosum ipsilaterally, due to corticotopic patterns of Ncam1 and other adhesion molecule expression (Figure 5). After reaching the midline, axons continue to co-fasciculate with axons matching their adhesion molecule levels, which now includes contralateral axons from symmetric cortical areas with the same expression gradients (Figure 6 and Movie S1). This callosal “handshake” would be a symmetric version of the proposed thalamocortical handshake^41,42^ shown to determine the topography and targeting of thalamocortical axons.^43^

In Ncam 1 gene deletion mice, the relationship between levels of potentially multiple contributing adhesion molecules in neighboring axons is not substantially changed compared to wild-type mice, since Ncam1 is uniformly deleted from all axons. This is predicted to result in somewhat lower overall levels of fasciculation, but not otherwise mismatched levels between neighboring axons– all axons would be regulated by interactions of their other homophilic molecules, but simply missing Ncam1.

In striking contrast, our genetic mosaic over-expression experiments markedly disrupt overall homophilic adhesion balance; targeted unilateral increase in Ncam1 levels in select axons interspersed with initially neighboring control axons creates a substantial mismatch between Ncam1 over-expressing and control ipsilateral neighbor axons, as well as with contralateral axons, which were not electroporated. This relative change in the most abundant homophilic adhesion molecule causes a significant mismatch that substantially alters the trajectory of targeted axons according to this levels-matching mechanism, resulting in high-Ncam1 expressing axons shifting their natural topographic trajectory toward target fields that endogenously express higher levels of Ncam1 (Figure 5).

Beyond the corpus callosum and interhemispheric connectivity, similar adhesion levels-matching mechanisms might be involved in the development of axon tract topography in non-symmetric systems. The observation of “pre-target sorting” and tract topography in many nervous system white matter tracts suggests such self-organizing mechanisms determining the positions of axons in a range of tracts via matching of graded adhesion levels between topographically neighboring axons.

This putatively ancestral mechanism of matched homophilic adhesion, which maintains ascending and descending axons in evolutionarily older systems organized as they grow, is revealed by our experiments to function similarly in the interhemispheric corpus callosum, the most recently evolved white matter projection, found only in eutherian mammals. Notably, this mechanism assumes an added element of axon guidance within the callosal system: after midline crossing, the callosum becomes a fully reciprocally symmetric system. Such a symmetric commissural tract naturally forms generally homotopic projections. Intriguingly, however, there are small subpopulations of non-homotopic callosal projections, such as those to the claustrum and peri-rhinal areas, and axons with mixed topography in the most ventral part of the callosum (Figures 1 and S1). These axons are predicted to actively modify their adhesion levels to avoid simply symmetric homophylic co-fasciculation, and/or to manipulate their timing so they meet symmetric partners outside the temporal window of homophilic adhesion levels matching, as we observed with heterochronic transplantation.

The work presented here poses new questions for investigation. Do elements of this mechanism of matching levels of homophilic - adhesion molecules generalize to development and topography of other axonal tracts, including other reciprocal projections? Are there other, related or unrelated molecular regulators participating in such axonal adhesion matching, and if so, do they include other hCAMs? Since the callosal system is fully symmetric and relatively evolutionarily recent, it is likely to include many, if not all, hCAMs present on axonal surfaces more generally, potentially ranging over wide concentration ranges, depending on potency and abundance. However, evolutionarily older and/or non-symmetric tracts might utilize a more limited set of hCAMs for such levels-matching mechanisms. There is high incidence of hCAM gene variants associated with human neurodevelopmental and neuropsychiatric conditions^44,45^, so identifying functions of these molecules in regulating precise topography of long-range connectivity in the brain might elucidate even subtly aberrant connectivity underlying these conditions.

## ACKNOWLEDGEMENTS

We thank Elise Caraker (Baltimore); Michelle Guo, Ioana Florea, Ben Noble, Thomas Addison, Hari Padmanabhan (Boston); and John Dumont (Athens) for technical support and useful discussions. We are grateful to the staff and instrumentation support of the following core facilities: the HSCRB Flow Cytometry Core, the Harvard FAS CSB Mass Spectrometry and Proteomics Resource Lab, and the UMSOM Center for Innovative Biomedical Resources through the Confocal Imaging Facility.

This work was supported by National Institutes of Health grants DP1 NS106665, R01 NS104055, and R21 NS104733 to JDM and DP2MH122398 to AP, as well as Autism Research Institute Autism Research Institute grant 30030461 to CB, with additional infrastructure support to JDM from the Max and Anne Wien Professor of Life Sciences fund. L.C.G. was partially supported by the Harvard Medical Scientist Training Program and NIH F31 NS080343. CB was additionally supported by the Schizophrenia & Psychosis-Related Disorders Training Grant of the Maryland Psychiatric Research Center T32MH067533.

## SUPPLEMENTARY MATERIAL

Figures S1-S6

Movie S1

## MATERIALS AND METHODS

### CONTACT FOR REAGENT AND RESOURCE SHARING

Further information and resources from this study are available upon request and availability. Requests should be directed to and will be fulfilled by the Lead Contacts, Jeffrey D. Macklis and Alexandros Poulopoulos.

### EXPERIMENTAL MODEL AND SUBJECT DETAILS

All animal experiments were designed and carried out in compliance with animal welfare guidelines of Harvard University and the University of Maryland. For information on mouse strains used, see Key Resources Table.

## METHOD DETAILS

### DNA constructs

Plasmid constructs were cloned with standard cloning methods and prepared for in utero injections using maxiprep plasmid kits (Qiagen). *Ncam1* Open Reading Frame (ORF) was subcloned from mouse brain cDNA corresponding to transcript NM_001113204.1 that translates into the 140 kDa isoform of NCAM1. Coding regions of all plasmids were verified by Sanger sequencing. See Key Resource Table for specific plasmid constructs.

### Neuroanatomical tracing

In vivo neuroanatomical tracing for mesoscale projection mapping was performed using nylon filters saturated with lipophilic dies in three colors. NeuroVue® Jade, Red, and Maroon Filter Squares (Polysciences) were cut into 1 mm^2^ pieces and placed subdurally through a craniotomy in P2 mice. Three squares of distinct colors were placed adjacent to one another along a rostrocaudal or mediolateral axis. Pups were deeply anesthetized and perfused 48h after labeling.

### Neuron micro-transplantation

Cortical neurons were prepared from micro-dissected lateral or medial regions of cortex from newborn mice. Neurons were prepared as described previously for neuron culture^46^ using papain digestion and trituration. Neurons were counted and equal numbers of GFP- and RFP-expressing neurons were combined in a cell suspension of approx. 50-100 cells/nl. The neuron suspension was loaded into a pulled glass micropipette attached to a digitally controlled nanojector (Drummond, Nanoject Variable). 50 nl of cell suspension was injected into the cortical parenchyma of dark littermate recipient mice, 50 μm deep to the dura, through a point craniotomy. Recipient brains in which neuronal cell bodies were positioned inappropriately beyond cortical grey matter into the callosal white matter were excluded from further analysis as a quality control step.

Micro-transplantation experiments were performed on litters from GFP +/- (transgene from mouse strain B6 ACTb-EGFP [JAX stock 003291]) and RFP +/- (transgene from mouse strain Ai9 [JAX stock 007909] females crossed with Vasa-Cre strain [JAX stock 006954] for constitutive RFP expression) back-bred onto an FVB background. These crosses resulted in GFP and RFP expression from constitutive β-actin promoters, with litters of mixed GFP, RFP, GFP/ RFP, and dark mice. Transplantations were performed only within littermates, thus using only litters with both green and red donor littermates, and dark recipient littermates. All experimental litters were within 6h of birth. A total of 6 dark recipient mice were analyzed at P7, each micro-transplanted with neurons prepared from 1-3 littermates of each color.

### In utero electroporation

In utero electroporations targeting layer II/III cortical neurons were performed at E15 as previously described.^23,47^ Briefly, plasmid DNA mixtures (5 μg/μl total DNA) were injected in one or both lateral ventricles, depending on the experiment, using a pulled and beveled glass micropipette. Five current pulses of 35 V were applied using a tweezer-trode or a tri-trode for light sheet experiments. Abdominal incisions were sutured. After term birth, electroporated mouse pups were screened for fluorescence using a fluorescence stereoscope (Leica MZ10 F with X-Cite Fire LED light engine).

### Fluorescent Small Particle Sorting (FSPS) of Growth Cones

Brains electroporated with GFP and RFP in opposite hemispheres, thus distinctly coloring lateral versus medial callosal projection neurons, were prepared for growth cone purification at P2. Growth cones were isolated and purified by FSPS as described previously.^23,48^ Briefly, 3-5 labeled brains were pooled immediately after dissection. The growth cone fraction was collected from an interface between 0.83 M and 2.5 M sucrose after homogenization under conditions of subcellular fractionation (isosmolar sucrose solution with low ionic strength) and buoyancy equilibration for 50 minutes under 250,000g in a vertical centrifugation rotor (VTi50, Beckman).

Fractions containing both red and green growth cones were diluted approximately 5-fold with PBS and freshly loaded into a sorter adjusted for sub-micron particle sorting (SORP FACSAriaII, BD instruments) to collect GFP+ and RFP+ growth cones. Particles were gated for size between 0.3 - 0.8 μm as determined by size-standard beads, and were collected based on single-color FACS plot gates for GFP or RFP fluorescence. 10^6^-10^7^ growth cones of each color were collected per experiment.

### Mass spectrometry

Purified growth cones were pelleted and subjected to on-pellet processing for label-free mass spectrometry, as described previously.^23,48^ Specifically, protein pellets were resuspended in 8 M urea, reduced with TCEP, alkylated with iodoacetamide, and digested with trypsin. Remaining material was sonicated i n acetonitrile, and digested further in trypsin.

Tryptic peptides were separated by a NanoAcquity UPLC pump (Waters) with a 5– 27% acetonitrile gradient in 0.1% formic acid. Peptides were ionized by electrospray and analyzed by tandem mass spectrometry (LC– MS/MS) on an LTQ Orbitrap Elite (Thermo Fischer). Sample order was randomized between replicate experiments. Data were analyzed by LFQ using MaxQuant and Perseus software.

### Immunolabeling and epifluorescence microscopy

Tissue for microscopy was obtained following intracardial perfusion of mice with PBS and 4% paraformaldehyde (PFA). Brains were sectioned coronally, parasagittally, or horizontally at 80 μm thickness on a vibrating microtome (Leica VT1000). Immunolabeling was performed in blocking solution with 5% bovine serum albumin and 0.1% Triton X-100. Primary and secondary antibodies were diluted 1:1000 in blocking solution and incubated with sections for at least 1h. Immunolabeled sections were mounted in Fluoromount-G Mounting Medium with DAPI (Thermo Fisher). For antibodies used, see Key Resources Table.

Epifluorescence imaging was performed on a Nikon 90i with automated stage controller and appropriate narrow bandpass filter sets. Whole brain section images were generated using EDF z-stack projections and image stitching through NIS-Elements (Nikon).

### In situ RNA hybridization

*In situ* hybridization was performed using an established protocol.^49,50^ Briefly, 50 μm thick cryosections mounted on glass slides (VWR, 48311-703) were hybridized with an appropriate probe labeled by digoxigenin, followed by incubation with anti-digoxigenin antibody conjugated to alkaline phosphatase. Sections were developed in a substrate solution containing 5-bromo-4-chloro-indolylphosphate and nitroblue tetrazolium chloride. The nucleotide region spanning coding and 3’ untranslated regions of *Ncam 1* cDNA (nucleotides 2675 to 3475, NM_001081445.2) was used as a probe, which is predicted to recognize Ncam1 isoforms 140 and 180.

### Tissue clearing and light sheet microscopy

Brains were cleared and imaged on a light sheet microscope as previously described^51^ based on a modified CUBIC protocol.^52^ Briefly, brains perfused with PFA at the indicated ages were immersed in CUBIC-L in a shaking water bath at 37 °C until white and grey matter were uniformly opaque. Then, brains were placed in Cubic-R+(M) at room temperature until transparent. Whole brains were imaged in Cubic-R+(M) on a Zeiss Light Sheet 7 and visualized with Arivis.

## QUANTIFICATION AND STATISTICAL ANALYSIS

Brain labeling and manipulation experiments were performed in triplicate. BEAM phenotypes were verified in multiple non-adjacent serial sections, and assessed visually based on separation of GFP from RFP axons.

Quantification of internally controlled micro-transplanation experiments was performed on N=6 recipient mice. Mice in which neuronal cell bodies were inadvertently transplanted into subcortical white matter were excluded from quantifications, as a quality control for appropriate grey matter transplantation. Labeled callosal axons were quantified in every-other section (non-adjacent), in 5 80 μm-thick coronal sections that included the entire mediolateral field. Each contralateral terminal axon segment was noted and represented, along with overlayed section outlines and transplantation injection sites as shown in Figure 2C and S3. The angles of each terminal axon segment from the intersection of the midline and a perpendicular line crossing the dorsal edge of the claustrum were measured and plotted on an angle histogram for each color (Figure 2C).

FSPS and subcellular proteomics were performed as two biological replicates, each from 3-6 pooled, bilaterally electroporated brains at P2. Mass spectrometry results were analyzed by MaxQuant using the MaxLFQ method for label-free quantification.^53^ Standard assignment of mass spectra was performed as previously with a false-discovery-rate cutoff of 0.05, and protein matching with a minimum of one unique peptide and FDR of 0.01 using the Andromeda search engine against version 83 of the Ensembl annotation. Raw LFQ values were averaged over the two replicates for medial and lateral growth cones.

Quantification of the phenotype from Ncam1 over-expression via the genetic mosaic BEAM system was performed in N=6 animals at P7 on matched coronal sections that contained the maximum number of labeled red and green axons. Fluorescence intensity was measured in each channel along a tangential line traversing layer VIb from the midline to the contralateral claustrum, as assessed by DAPI staining. Labeled callosal axons intersecting this line were exiting the callosal white matter to innervate the contralateral grey matter. The cumulative intensity in each channel along this line was normalized to 1. The line was divided into 50 equal bins, and the value of the integral intensity was measured for each channel, expressed as a proportion of the total cumulative intensity. This yielded the fractional normalized intensities for each channel in each bin. These values correspond to the fraction of total axons exiting the callosal white matter at the bin position for each mosaic condition (Ncam1 over-expression or interspersed control within the same area of the same mouse). These values were plotted on cumulative histograms for the 50 bins (Figure 5B).

## KEY RESOURCES TABLE

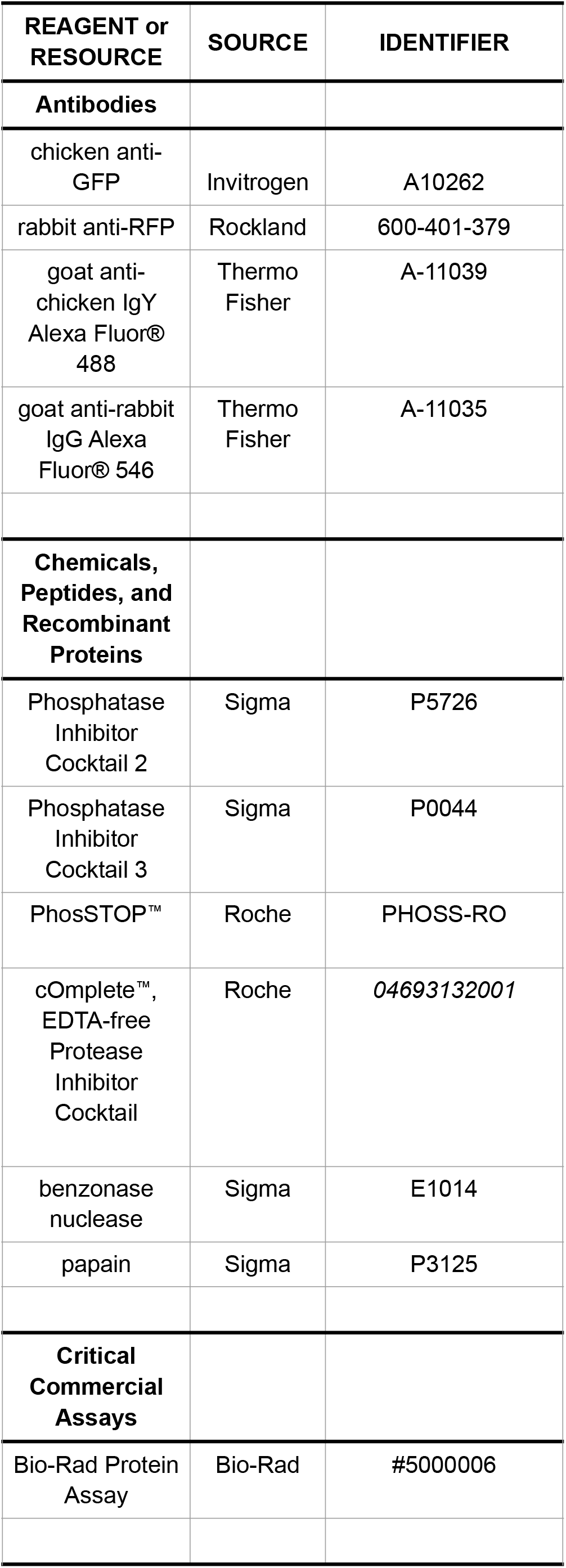
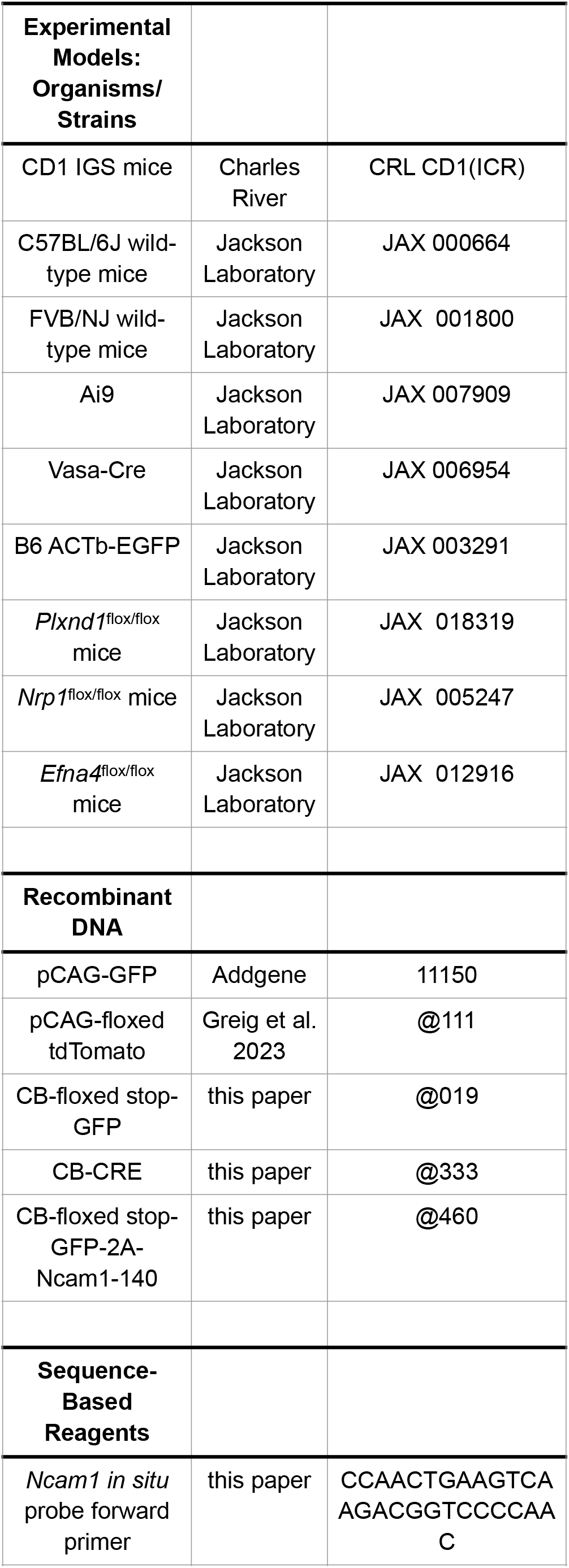
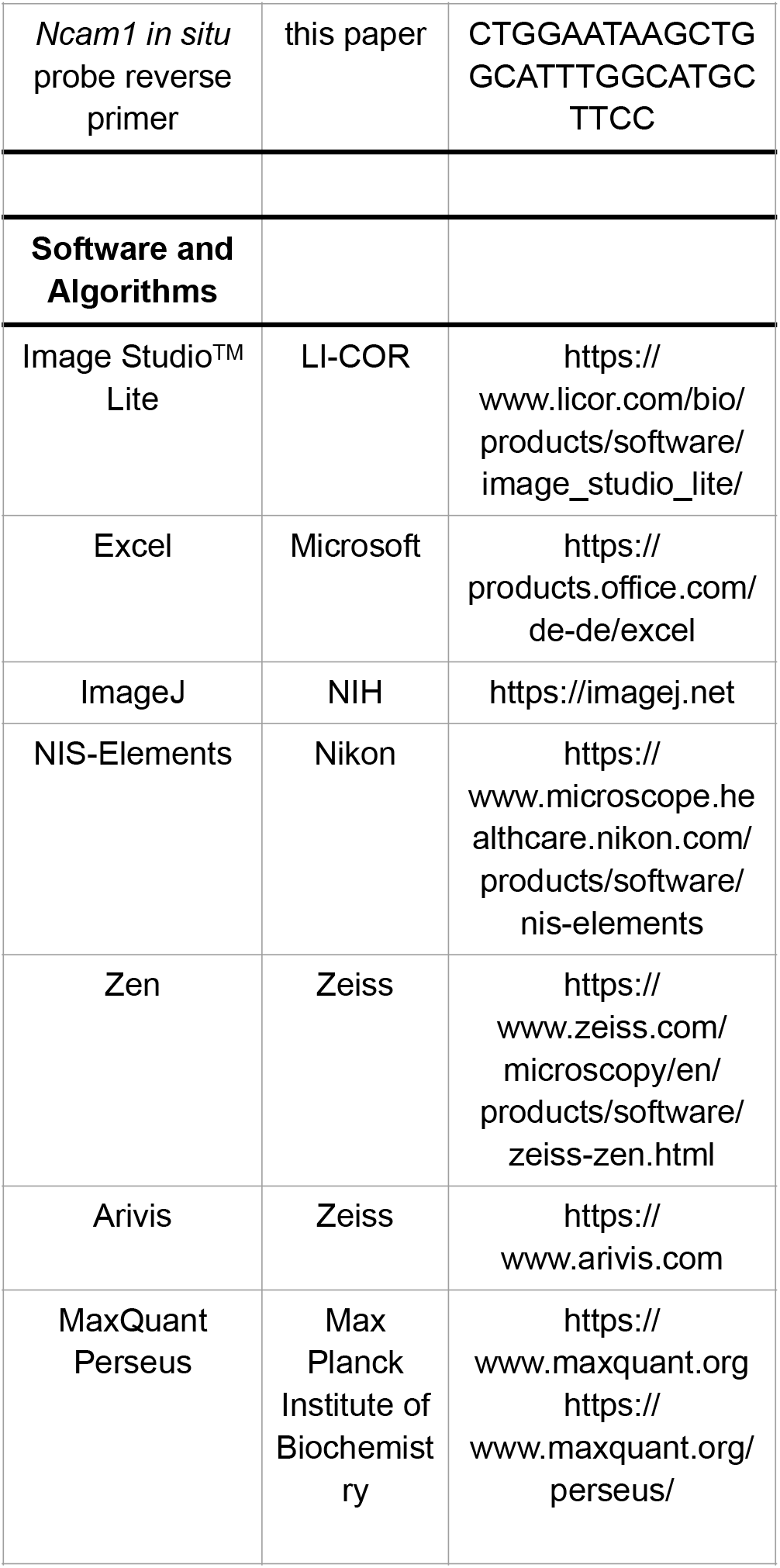

