## Supplementary material for "Symmetry in levels of axon-axon homophilic adhesion establishes topography in the corpus callosum and development of connectivity between brain hemispheres": Figures S1-S6

#### Figure S1

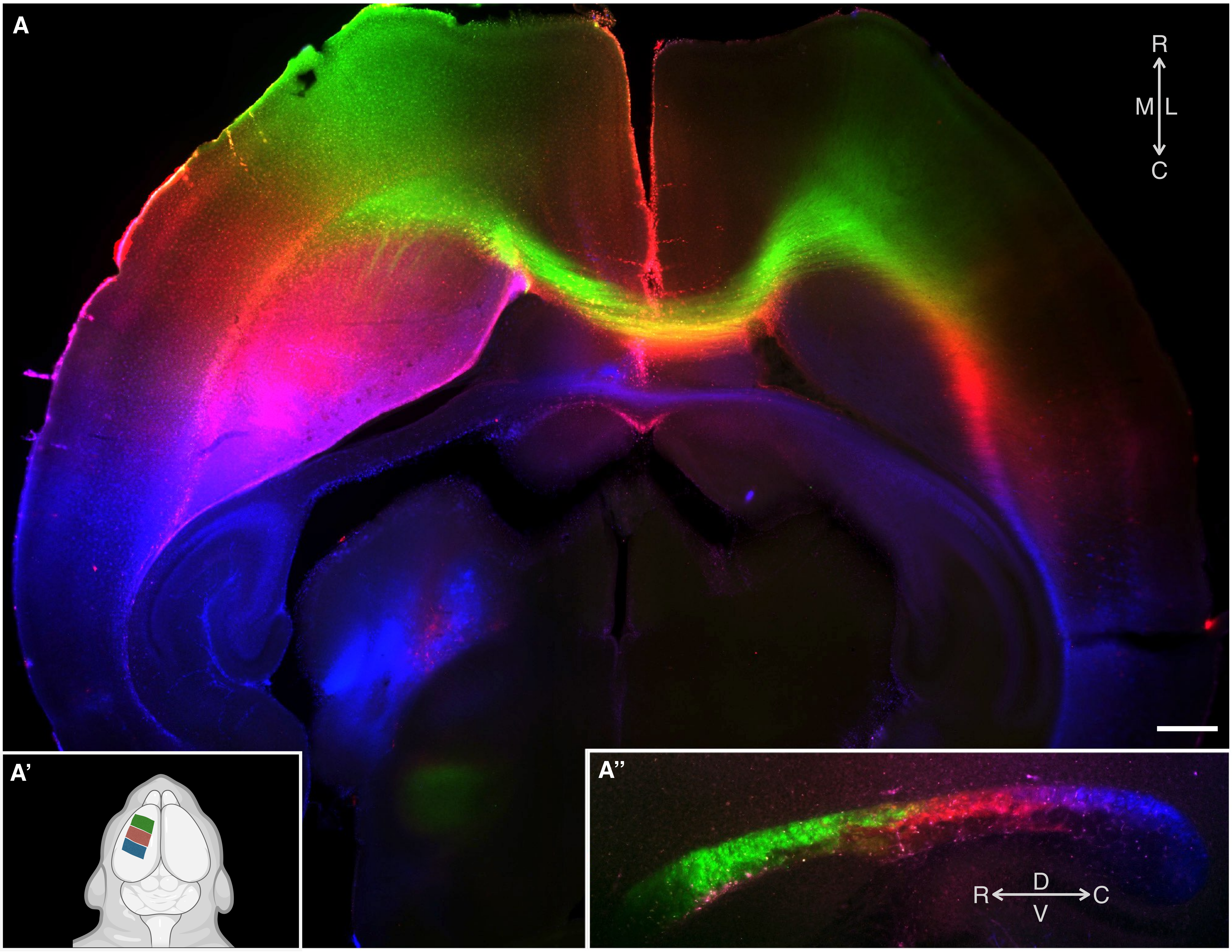


**Figure S1. Mesoscale topographic mapping of cortical projections along the rostrocaudal corticotopic axis**
Neuroanatomical tracers in three colors placed via small saturated filter paper squares on the cortical surface along orthogonal topographic axes (rostrocaudal here, mediolateral in Figure 1) in infant mice reveal the internal topography along white matter tracts. (A) Horizontal section of P4 brain labeled along the rostrocaudal axis, schematized in (A'). (A") Sagittal view of the corpus callosum at the level of the midline. Callosal axon projections (projecting orthogonal to the page) display corticotopic topography throughout the callosal white matter tract such that the rostrocaudal axis in the cortex is maintained as the rostrocaudal axis in the callosum.

### Figure S2

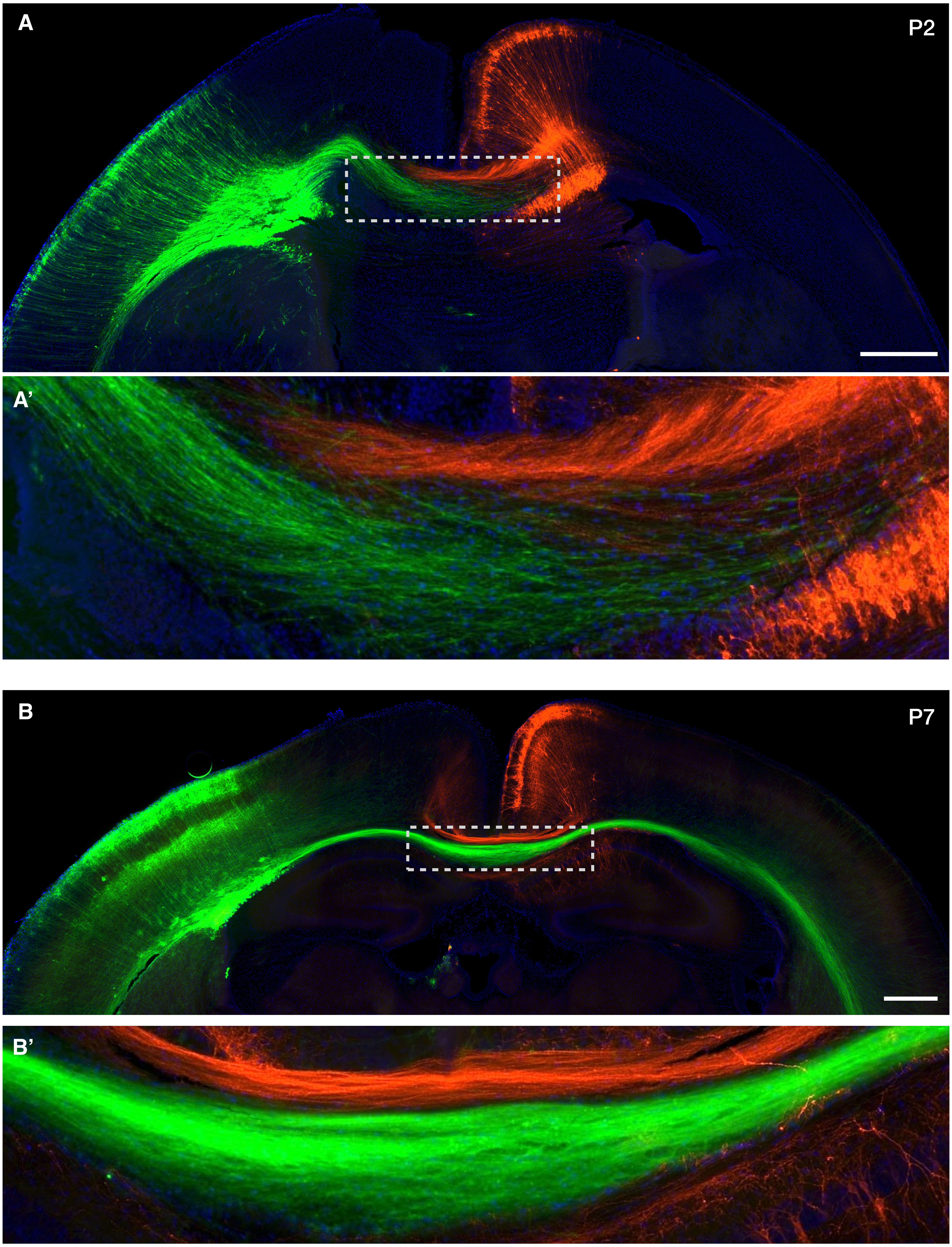


**Figure S2. *In utero* labeling of medial and lateral callosal projection neurons in development shows early callosal projection topography**

(A) Coronal section of P2 mouse brain following asymmetric, dual-hemisphere *in utero* electroporation to label developing medial (red) and lateral (green) superficial layer cortical projection neurons in apposing hemispheres. (A’) Inset of dashed box in A reveals red and green axons from contralateral hemispheres sharply segregating once they encounter each other crossing the midline. (B) P7 coronal section of mouse brain following the same type of asymmetric, dual-hemisphere electroporation. (B’) Inset of dashed box in B, reinforcing the consolidation of sharp segregation of medial red and lateral green callosal axons avoiding contact over long distances with opposite trajectories within the axon tract.

**Figure S3**

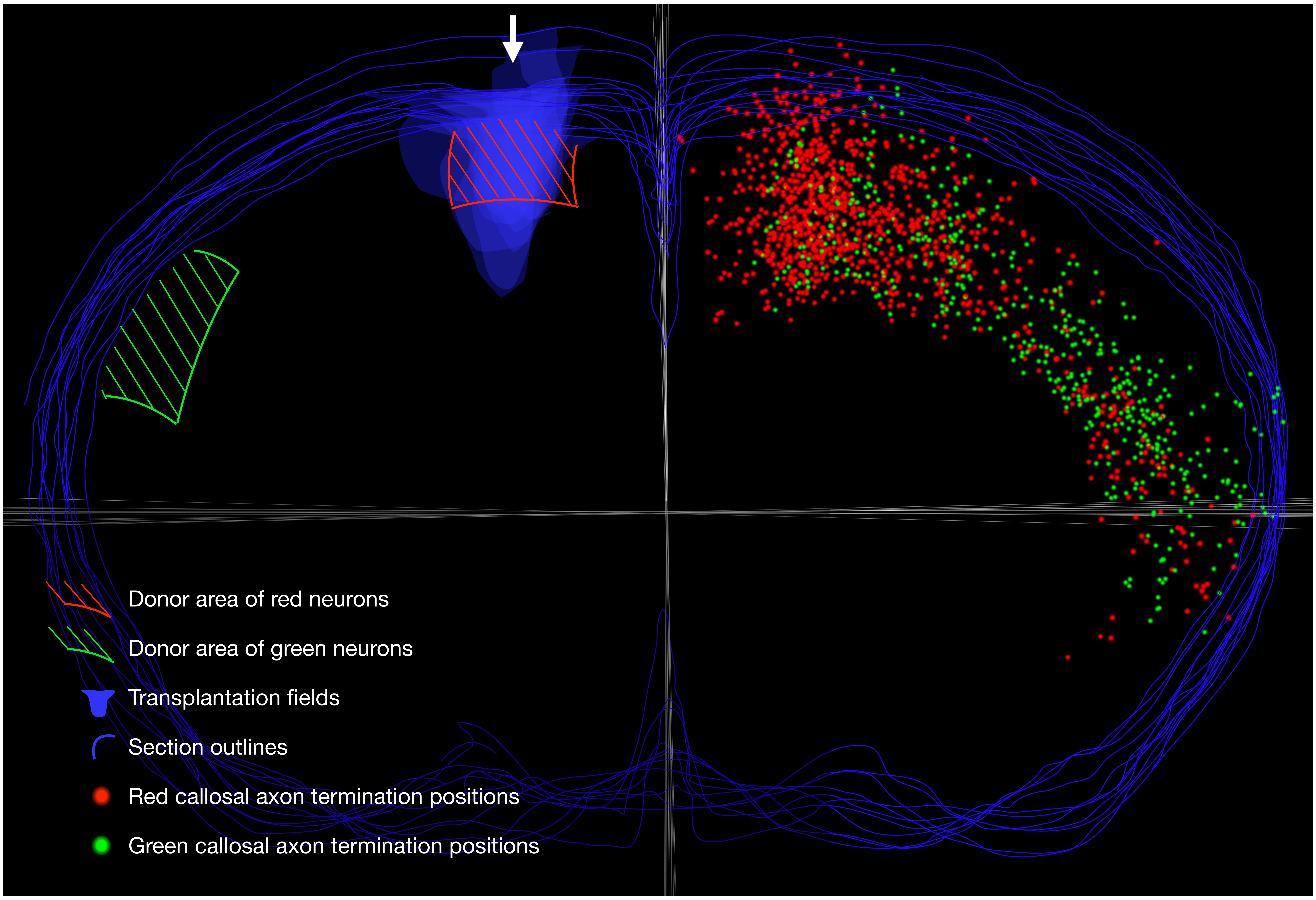


**Figure S3. Heterotopic micro-transplantation of lateral and medial cortical neurons reveals intrinsic callosal axon targeting**
Data aggregation of heterotopic micro-transplantation experiment shown in Figure 2. Both ectopic transplanted cells from lateral cortex (schematized as green shaded region) and internal-control transplanted cells from medial cortex (schematized as red shaded region) were together micro-injected into medial cortex (arrow) of 6 dark recipients. Schematic overlays 3 of the 5 traced sections from each of the 6 brains quantified in Figure 2C. Blue outlines coronal section contour and highly reproducible injection transplantation fields in ipsilateral medial cortex. Red and green dots indicate corresponding axon termination positions in contralateral gray matter. Grey lines indicate midline and meridian axes used to quantify axon position as degrees from midline at intersection.

### Figure S4

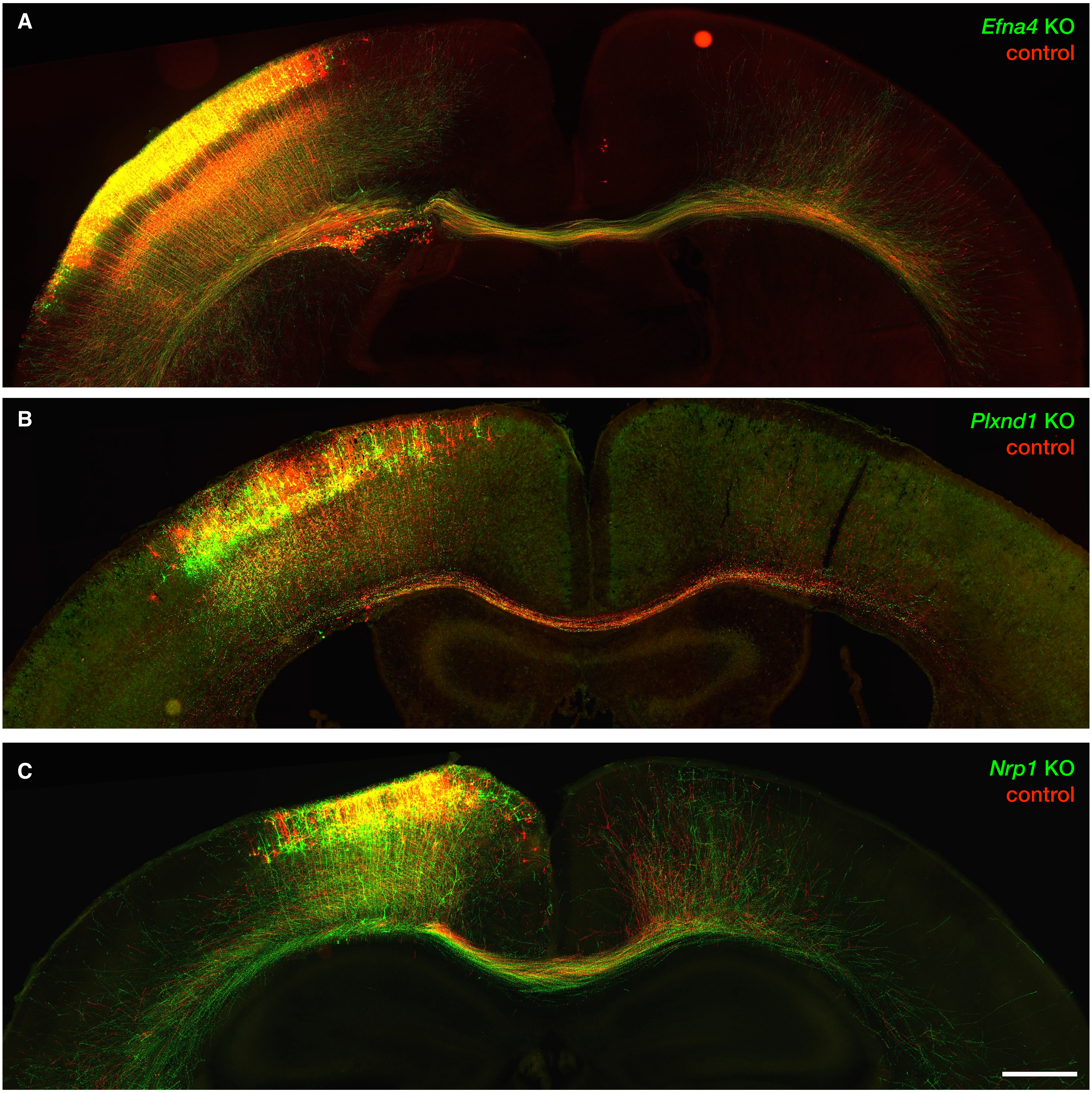


**Figure S4. *In utero* mosaic gene deletion with spatially co-localized, interspersed internal control neurons via BEAM[26] reveals that the canonical axon guidance molecules Ephrin-A4, Plexin-D1, and Neuropilin-1 are dispensable for early callosal projection topography**

Genetic mosaic (via BEAM[26]) *in utero* electroporation of superficial layer callosal projection neurons stochastically expressing either green, Cre-delivering or red, control plasmids to interspersed neurons within the same electroporation field. Green "experimental" neurons express GFP together with Cre to delete floxed alleles. Red "control" neurons express RFP alone, and serve as region- and subtype-specific internal controls, and exhibit normal projection targeting. Coronal sections of P7 brains electroporated at E15.5 in homozygous (A) *Efna4*$^{flox/flox}$, (B) *Plxnd1*$^{flox/flox}$, and (C) *Nrp1*$^{flox/flox}$ mice. Neurons with deletion of Ephrin-A4, Plexin-D1, or Neuropilin-1 (green) have unaltered, normal contralateral targeting of callosal axons compared to interspersed wild-type control neurons (red).

### Figure S5

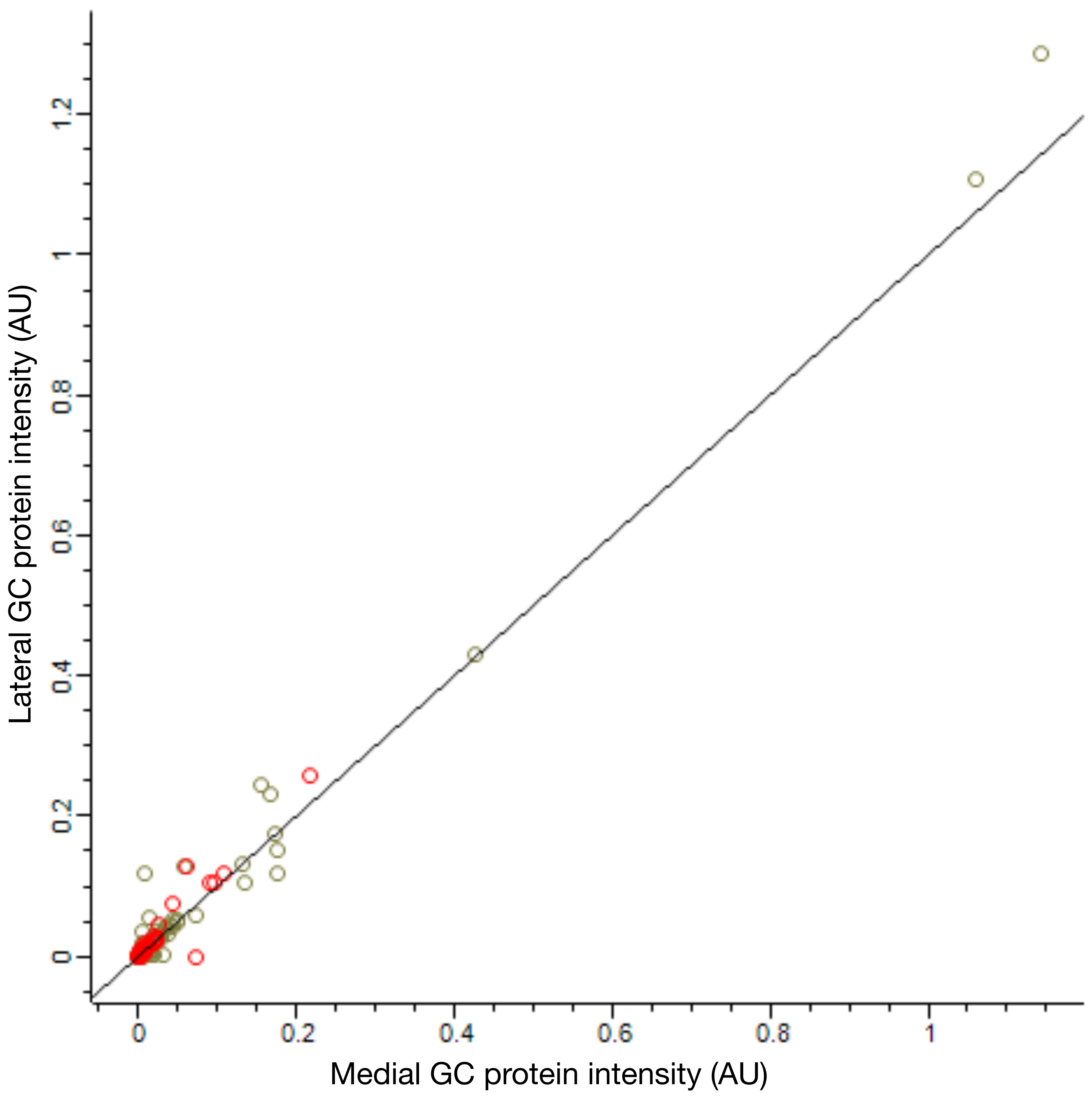


**Figure S5. Comparative subcellular proteomics of medial versus lateral callosal growth cones**

Scatter plot extension of Figure 4C showing protein amounts in medial vs. lateral callosal neuron growth cones, as quantified by MaxLFQ. This is the full quantitative range scatter plot of Figure 4C, including the high-abundance points beyond the linear range. Red circles represent membrane proteins; grey circles represent non-membrane proteins. Each circle represents the average protein intensity measurements from medial (x-axis) and lateral (y-axis) growth cones from two biological replicates, each from 3-5 pooled dual-electroporated brains.

#### Figure S6

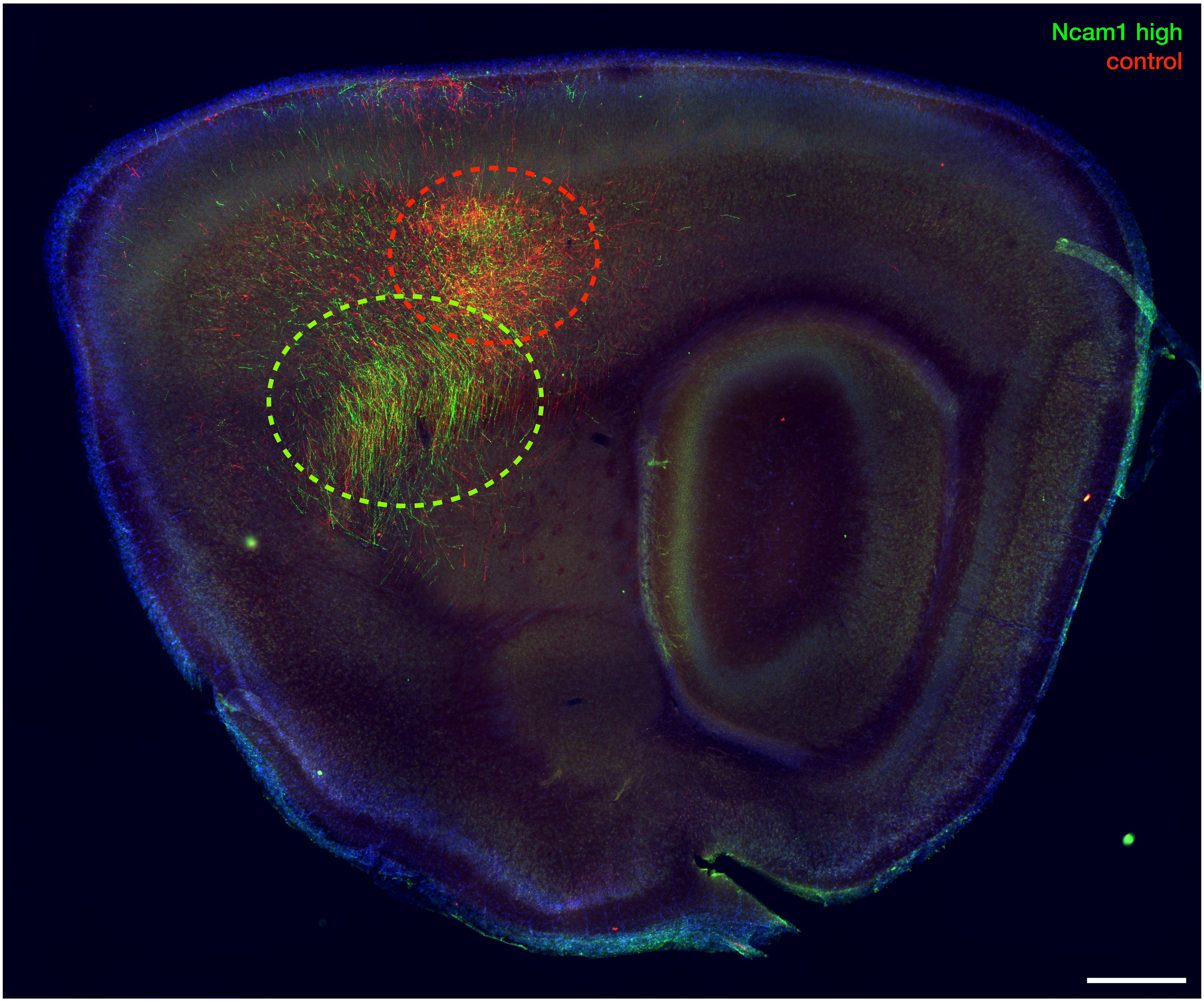


**Figure S6. Ncam1 expression levels determine contralateral axon target fields of callosal projection neurons**

Image of a contralateral parasagittal section from the experiment shown in Figure 5A. Following genetic mosaic *in utero* electroporation of superficial layer callosal projection neurons via BEAM[26], green electroporated neurons over-express Ncam1, while red electroporated neurons remain wild-type control. A large fraction of Ncam1-overexpressing axons (green) project to a non-canonical ventral target field (green dashed ellipse) in the contralateral grey matter, compared to the normal, more dorsal contralateral target field of wild-type (red) control axons (red dashed ellipse).
